# Epigenetic downregulation of HER2 during EMT leads to tumor resistance to HER2-targeted therapies in breast cancer

**DOI:** 10.1101/2020.03.24.006379

**Authors:** Babak Nami, Avrin Ghanaeian, Zhixiang Wang

## Abstract

HER2 receptor tyrosine kinase (encoded by *ERBB2* gene) is overexpressed in approximately 25% of all breast cancer tumors (known as HER2-positive breast cancers). Overexpression of HER2 causes overactivation of downstream receptor tyrosine kinase pathways including PI3K/Akt and MAPK pathways and is a poor prognosis factor in breast cancer. Tyrosine kinase inhibitor lapatinib and anti-HER2 monoclonal antibodies trastuzumab and pertuzumab are FDA-approved HER2-targeted drugs for treatment of HER2-positive breast cancers. However, development of de novo resistance to HER2 blockade occurs in majority of patients after treatment started. Resistance to HER2 targeting therapies partially due to the loss of HER2 expression on their tumor cells during the treatment. But little is known about the exact mechanism of loss of HER2 on originally HER2-positive tumor cells. Downregulation of extracellular HER2 by metalloproteinases during epithelial-mesenchymal transition (EMT) in trastuzumab-resistant/lapatinib-sensitive cells has been shown by limited studies, however, the mechanism of ERBB2 gene silencing during EMT and in the mesenchymal-like cells derived from trastuzumab-resistant/lapatinib-resistant HER2-positive breast tumors was entirely unknown. In this study, hypothesized that EMT abrogates HER2 expression by chromatin-based epigenetic silencing of *ERBB2* gene as a mechanism of acquired resistance to HER2-targeted therapies. we found that HER2 expression is positively and negatively correlated with the expression of epithelial and mesenchymal phenotype marker genes respectively in breast cancer tumors. We also found that chromatin of *ERBB2* gene in HER2-high epithelial-like breast cancer cells is active, while, the chromatin is inactive in HER2-low mesenchymal-like cells. HER2-low breast cancer cell line also revealed less promoter-enhancer interaction and small chromatin loops compared to the HER2-high cell lines. The lower HER2 expression, the higher EMT phenotype, and inactivated chromatin all were found correlated with a lower response to lapatinib. The higher EMT phenotype was found correlated with a lower response to lapatinib. We also found that induction of EMT of HER2-positive breast cancer BT474 cells results in downregulated HER2 expression and lower binding rate of trastuzumab to the cells. These results show that the downregulation of HER2 in mesenchymal-like cells in the culture of HER2-positive breast cancer cell lines was due to *ERBB2* gene silencing by epigenetic reprogramming of the cells during EMT. These results indicate that *ERBB2* gene silencing by epigenetic regulation during EMT is the main mechanism of resistance of HER2-positive breast cancer cells to trastuzumab and lapatinib.

## INTRODUCTION

HER2 receptor tyrosine kinase (encoded by *ERBB2* gene) is a 1255 amino acid transmembrane epidermal growth factor receptor tyrosine kinase protein with 185 kDa molecular weight [1,2]. HER2 contains four extracellular domains (ECD; domains I-IV; amino acids 1-641), an extracellular juxtamembrane region (EJM; amino acids 642-652) a transmembrane domain (TM; amino acids 653-675), an intracellular juxtamembrane region (IJM; amino acids 676-730), an intracellular tyrosine kinase domain (TK; amino acids 731-906), and a C-terminal tail (amino acids 907-1255) [3–6]. The ECD of HER2, by turn, consists of four subdomains including two leucine-rich domains (domains I and III) and two cysteine-rich domains (domains II and IV) containing receptor dimerization motif [3–6]. HER2 does not need ligand to be activated and can make dimer with another HER2 (homodimerization) or other HER family receptors (HER1/EGFR and HER3; hetero-dimerization). HER2 dimerization causes activation of its kinase domain which phosphorylates downstream mediators and promotes the receptor tyrosine kinase signaling cascades (PI3K/Akt, PLC-γ and MAPK pathways).

Overexpression of HER2 protein occurs in 20-30% of all breast cancer tumors (known as HER2-positive breast cancers) majority of it due to *ERBB2* gene amplification [2,7–9]. HER2 overexpression results in overactivation of downstream PI3K/Akt, PLC-γ and MAPK pathways leading to increased tumor cell growth, survival, motility, and invasion [10]. Targeting HER2 by small molecule inhibitors and monoclonal antibodies is the current therapy for HER2-positive breast cancer that outcome significant tumor regression in the patients [11,12]. Lapatinib, a small molecule dual inhibitor of tyrosine kinase activity of HER2 and EGFR, and trastuzumab (Herceptin) and pertuzumab (Perjeta) which are anti-HER2 humanized monoclonal antibodies targeting ECD of HER2 approved by FDA to treat patients with early-stage and metastatic HER2-positive breast cancer as an adjuvant in combination with taxane therapy [11,13,14].

Unfortunately, development of acquired resistance to HER2-targeted therapies continues as a big obstacle in the treatment of HER2-positive breast cancer. Approximately 60-70% of HER2-positive breast cancer patients develop de novo resistance to trastuzumab, partially due to the loss of HER2 expression on their tumor cells during the treatment [11,12,15]. A suggested mechanism of trastuzumab resistance is cleavage and shedding of HER2. Cleavage from EJM region will result in shedding of the extracellular part of HER2 and production of p95HER2 still encored at the plasma membrane. While, in the case of cleavage from IJM region, the intracellular part of HER2 will shed and the extracellular part will remain at the plasma membrane [11,12,16,17]. Both cleavage cases that can take place by proteinases during epithelial-mesenchymal transition (EMT) of HER2-positive breast cancer, that leads to emergence of tumor cells resistant to trastuzumab but still sensitive to lapatinib [12]. Notably, little is known about the exact mechanism of lapatinib resistance in HER2-positive breast cancer.

We previously reviewed our hypothesis that suggests role of EMT in development of de novo resistance to HER2-targeted therapies [12]. Generally, epithelial-like cells highly express HER2, whereas mesenchymal cells are majorly HER2-negative or HER2-low. This shows that mesenchymal-like cells show resistance to trastuzumab, suggesting that trastuzumab-resistance may link to EMT. JIMT-1 cell line is HER2-positive breast cancer cells that quickly develop resistance to HER2 [18]. A study showed that JIMT-1 was composed of approximately 10% CD44+/CD24-breast cancer stem cells in initial cultures, however, this level rose to 85% at the late-passages [19]. Concurrently, the level of HER2 expression significantly reduced in late-passage cultures when compared to the early cultures which was associated with the development of trastuzumab-resistance [19]. This phenomenon may explain the resistance of HER2-high breast tumors to trastuzumab due to an increased population of HER2-low CD44+/CD24 mesenchymal cells at the late-passages. Further, the CD44+/CD24-cells escape from trastuzumab-mediated antibody dependent cellular cytotoxicity (ADCC) and survive the immunoselection process in breast cancer cells co-cultured with natural killer (NK) cells and trastuzumab [20]. This resistance may be attributed to the reduced HER2 expression levels on their surface [20]. These pieces of evidence show that downregulated HER2 and therefore decreased response to trastuzumab are parts of the intrinsic regulation of mesenchymal breast cancer cells. However, the mechanism of this regulation is not yet studied.

Here hypothesize that wide-scale epigenetic reprogramming during EMT could be the mechanism of *ERBB2* gene silencing and development of resistance to HER2-targeted agents. The aim of this study was to study the chromatin-based epigenetic mechanism of *ERBB2* gene silencing during EMT and its relation to anti-HER2 therapy resistance.

## RESULTS

### Mesenchymal breast cancer cells show lower *ERBB2* gene expression

To investigate whether expression level of *ERBB2* gene is correlated with the expression of EMT marker genes, we analyzed the RNA-seq expression of *ERBB2* gene, 12 epithelial marker genes *(ELCAM, CD24, CDH1, F11R, FOXA1, KRT7, KRT8, KRT18, KRT19, MUC, NECTIN2, NECTIN4)* as well as 12 mesenchymal marker genes (*CD44*, *CTNNB1, FOXC1, MYC, NOTCH1, NOTCH2, SNAI2, SOX10, TWIST2, VIM, ZEB1*, *ZEB2)* in 1,904 breast cancer tumor samples studied by METABRIC study [21]. We used cBioPortal portal [22] to investigate correlations between the mRNA levels of *ERBB2* and the EMT markers in each tumor sample. The result showed a significant positive correlation between the expression of *ERBB2* and all the epithelial marker genes (Figure 1) and a negative correlation between ERBB2 and all the mesenchymal marker genes (Figure 2).

**Figure 1.**
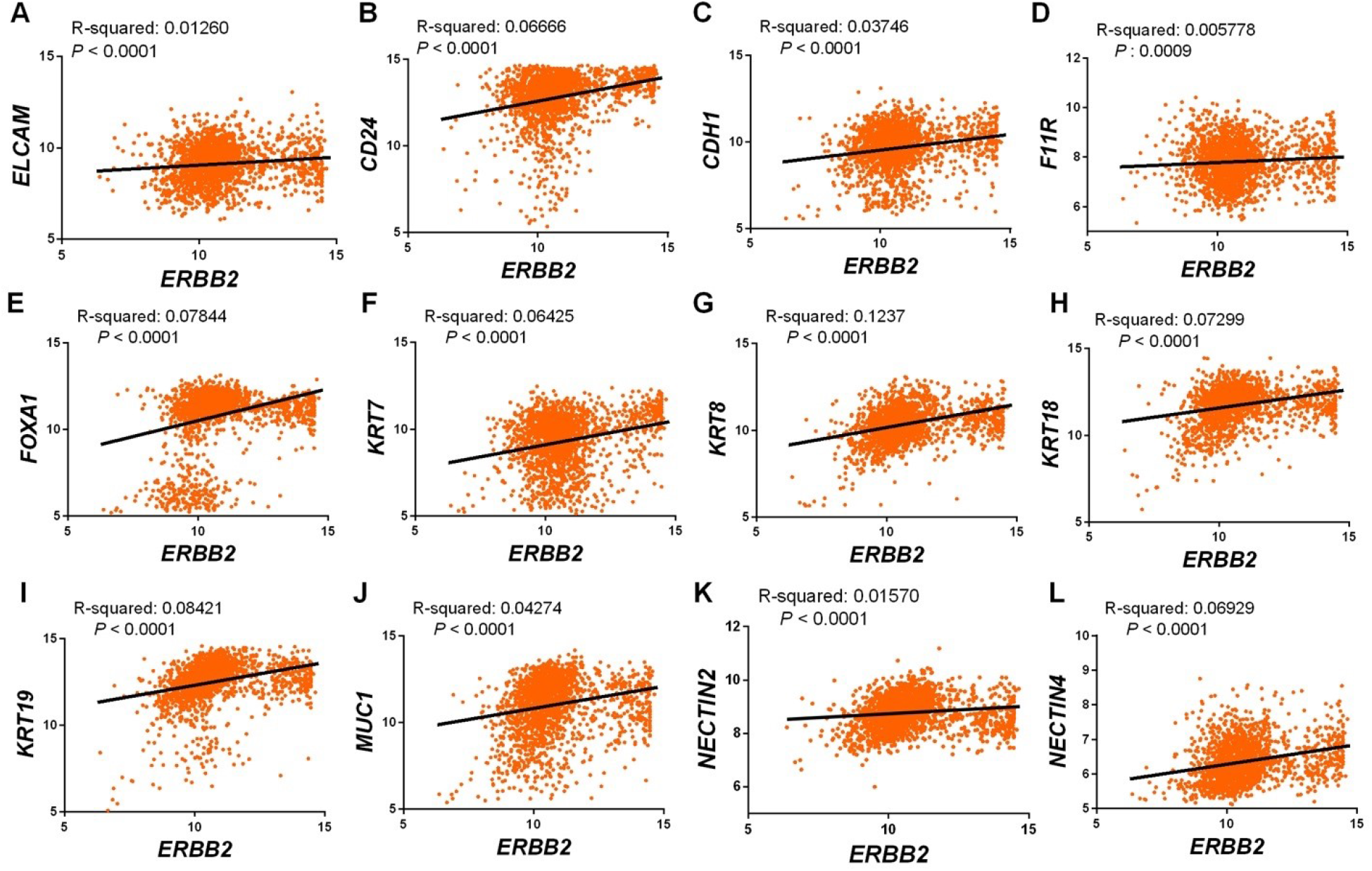
Correlation between mRNA expression of *ERBB2* and epithelial-like cell markers in breast cancer tumors. The expression value is presented by Z-score fold changes RNA-seq expression (v2 RSEM). Data source: normalized RNA-seq data from 1,904 breast tumors studied by METABRIC study [21] and available from cBioPortal portal [22] available at https://cbioportal.org.

**Figure 2.**
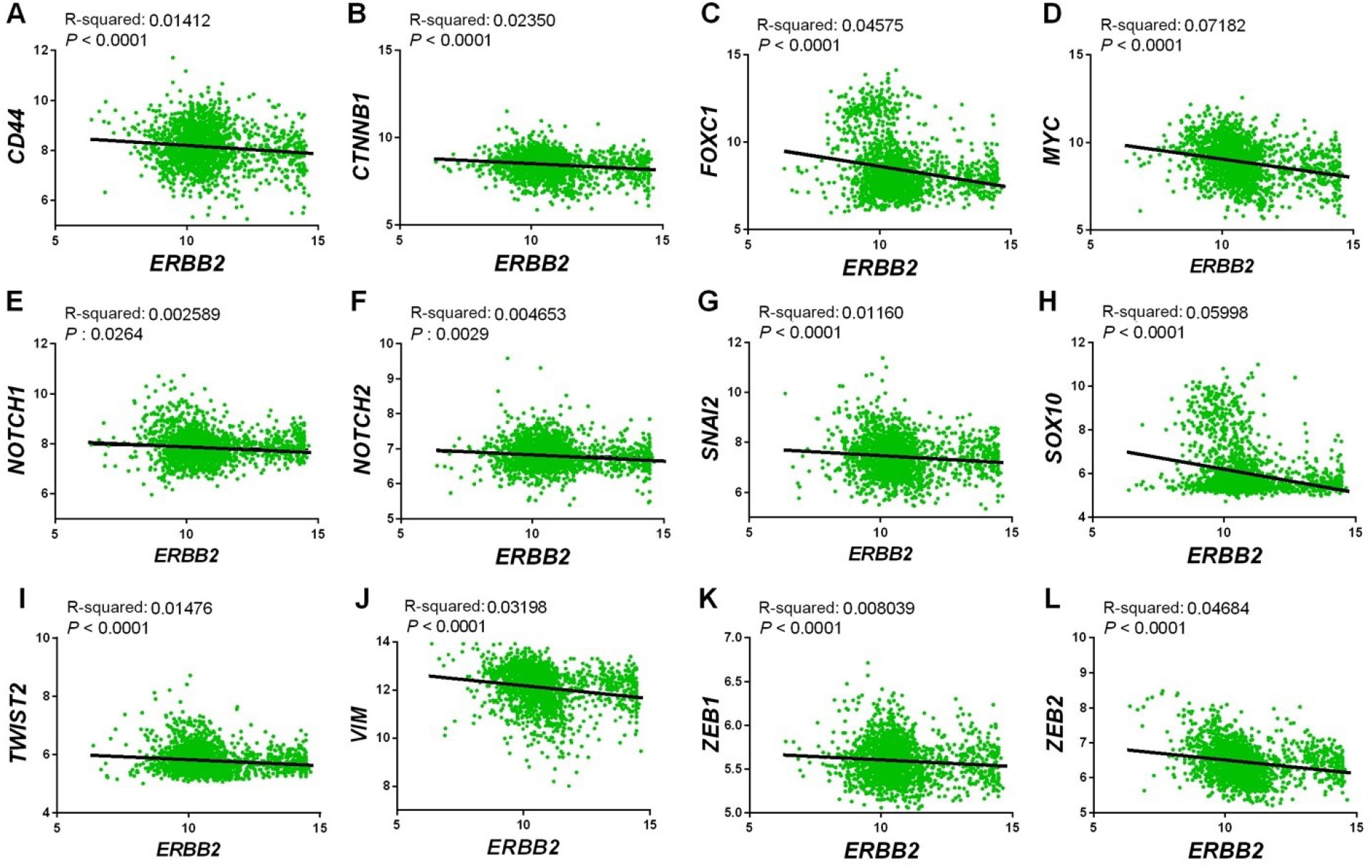
Correlation between mRNA expression of *ERBB2* and mesenchymal-like cell markers in breast cancer tumors. The expression value is presented by Z-score fold changes RNA-seq expression (v2 RSEM). Data source: normalized RNA-seq data from 1,904 breast tumors studied by METABRIC study [21] and available from cBioPortal portal [22] available at https://cbioportal.org.

We also analyzed the expression of *ERBB2,* epithelial marker gene *MUC1,* mesenchymal marker gene *VIM,* and GAPDH in 38 breast cancer cell lines (AU565, BT-20, BT474, BT-549, CAL-51, CAMA-1, DU4475, HCC-1143, HCC-1187, HCC-1395, HCC-1419, HCC-1428, HCC-1500, HCC-1569, HCC-1599, HCC-1806, HCC-1937, HCC-1954, HCC-202, HCC-2218, HCC-3, HCC-38, HCC-70, Hs578T, MCF7, MDA-MB-175-VII, MDA-MB-231, MDA-MB-361, MDA-MB-436, MDA-MB-453, MDA-MB-468, SKBR3, T47D, UACC-812, UACC-893, ZR-75-1, ZR-75-30, ZRT) to study the correlation between mRNA expression levels of *ERBB2* and EMT marker genes. For this, two normalized microarray expression datasets (GEO accession numbers: GSE50811 [23] and GSE66071 [24]) available from NCBI GEO (Gene Expression Omnibus) database were analyzed. As shown in Figure 3, comparatively, the expression of *ERBB2* was positively and negatively correlated with the expression of respectively *MUC1* and *VIM* in most of the cell lines.

**Figure 3.**
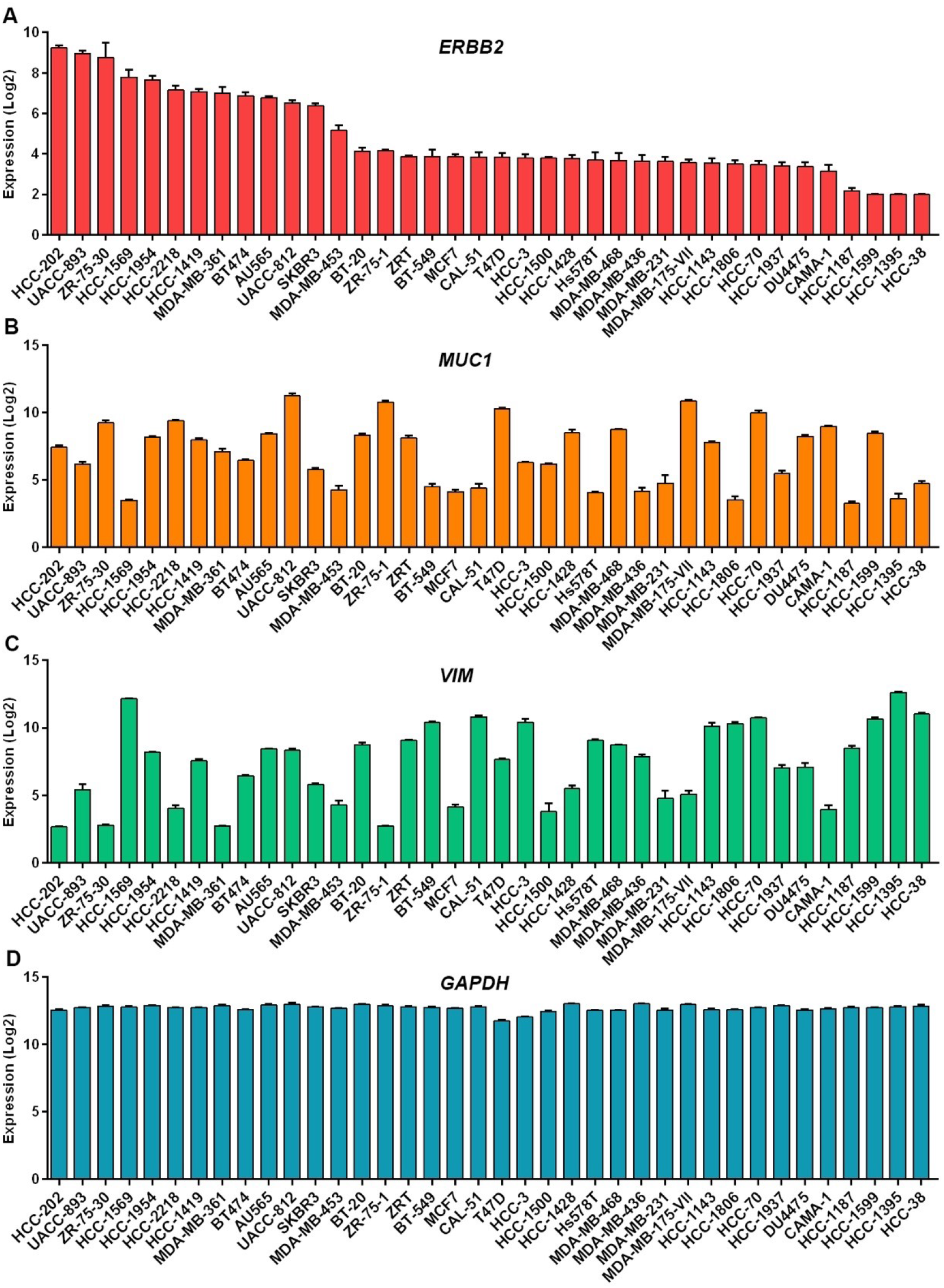
mRNA expression levels of *ERBB2, MUC1, VIM* and *GAPDH* in breast cancer cell lines. Data source: Microarray expression profiling (GEO accession numbers: GSE50811 [23] and GSE66071 [24] available from GEO database.

These results show positive coloration between *ERBB2* gene expression and epithelial phenotype, and a negative correlation between *ERBB2* gene expression and mesenchymal phenotype in breast cancer. This suggests that the expression of *ERBB2* gene in epithelial-like breast cancer cells is higher than that in mesenchymal-like breast cancer cells, suggesting that mesenchymal cells mostly show low *ERBB2* gene expression compared to epithelial-like breast cancer cells.

### Promoter CpG islands methylation signature of *ERBB2* in epithelial-like and mesenchymal-like breast cancer cells

Transcription of genes is highly under control of promoter epigenetic regulation in DNA (CpG island methylation) and chromatin (histone protein modification) levels. To study the mechanism of low *ERBB2* gene expression we investigated promoter CpG island methylation signature of *ERBB2* gene in breast cancer cell lines with high *ERBB2* expression (BT474, HCC-1954, MDA-MB-453, SKBR3), and those with low *ERBB2* expression (BT20, MCF7, MDA-MB-231, MDA-MB-468, SUM-159PT, T47D). Array expression and genome tiling array methylation data of the cells were obtained from GEO database (Accession number: GSE44838 [25])

The mRNA expression levels of *ERBB2* in the cells are shown in Figure 4A. The result showed a positive correlation between the expression of *ERBB2* and *FOXA1* (epithelial-like cell marker), and the negative correlation between *ERBB2* expression and the expression of *FOXC1* (mesenchymal-like cell marker) in all cell lines except HCC-1954 (Figure 4A-C). Despite the different *ERBB2* expression levels of the cell lines, no significant difference was found between the cell lines in terms of the CpG island methylation (Figure 4D). These results show that low *ERBB2* expression levels in the mesenchymal-like cells are not due to promoter CpG island methylation.

**Figure 4.**
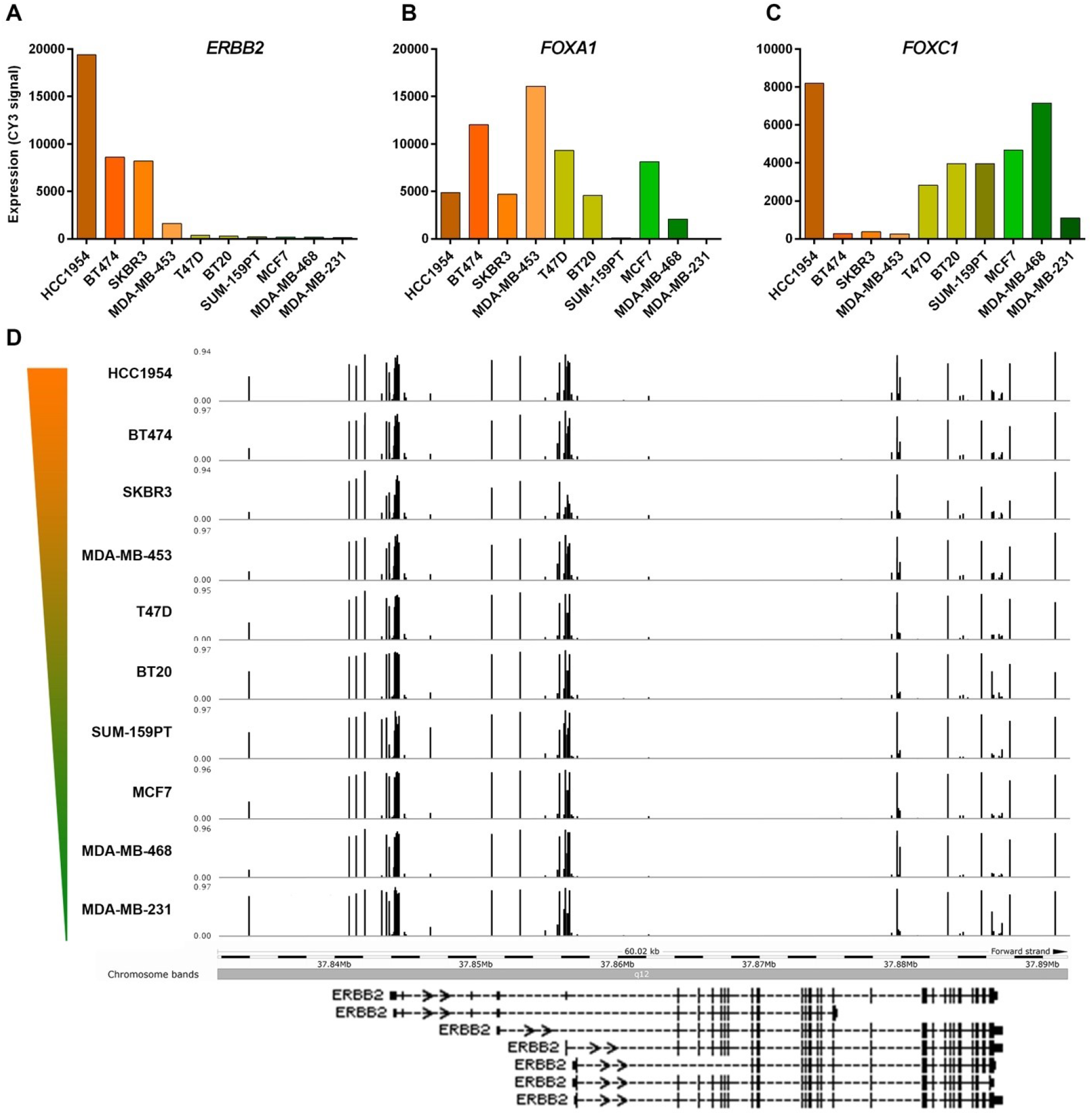
CpG island methylation profiles of *ERBB2* promoter in 10 breast cancer cell lines. (**A**) mRNA expression levels of *ERBB2, FOXA1,* and *FOXC1* genes in breast cancers. (**B**) Methylation levels of promoter CpG islands in the cell lines. Data source: Array expression profiling and genome tiling array methylation profiling (GEO Accession number: GSE44838 [25]) available from GEO database. The color gradient bar indicates HER2 expression level. Genomic coordinate: chr17:37,834,978-37,897,500 (GRCh37/hg19 assembly).

### EMT regulator transcription factors bind to *ERBB2* regulatory elements

Since DNA level epigenetic regulation (promoter CpG island methylation) has no role in the expression of *ERBB2,* we hypothesized that chromatin level epigenetic regulation (promoter and enhancer histone protein modification) may control different levels of *ERBB2* expression in the breast cancer cells. To examine this, we first studied transcription factors that directly bind to *ERBB2* chromatin identified by the ChIP-seq indexed in ChIPbase v2.0 database. Result identified enrichment of a totally 82 transcription factors at the region of 10 kbp upstream and 10 kbp downstream of *ERBB2* gene motif Y in 3,740 human biological samples (Figure 5A). Of 82 transcription factors 8 (CDX2, FOXA1, FOXA2, KLF9, MBD3, MXI1, RUNX3, SP1) were epithelial status maintenance regulators, and 31 (ATF2, E2F1, E2F6, E2F7, EGR1, ELF2, ETS1, ETV1, FOS, FOXM1, FOXP1, FOXP2, GATA1, GATA2, GATA3, GATA6, HOXC9, JUNB, JUND, KDM5A, MAX, MAZ, MYC, MZF1, NANOG, RELA, SMAD4, STAT4, STAT5A, TEAD6, ZBTB7A) were EMT inducers which are master regulators of mesenchymal status maintenance (Figure 5B).

**Figure 5.**
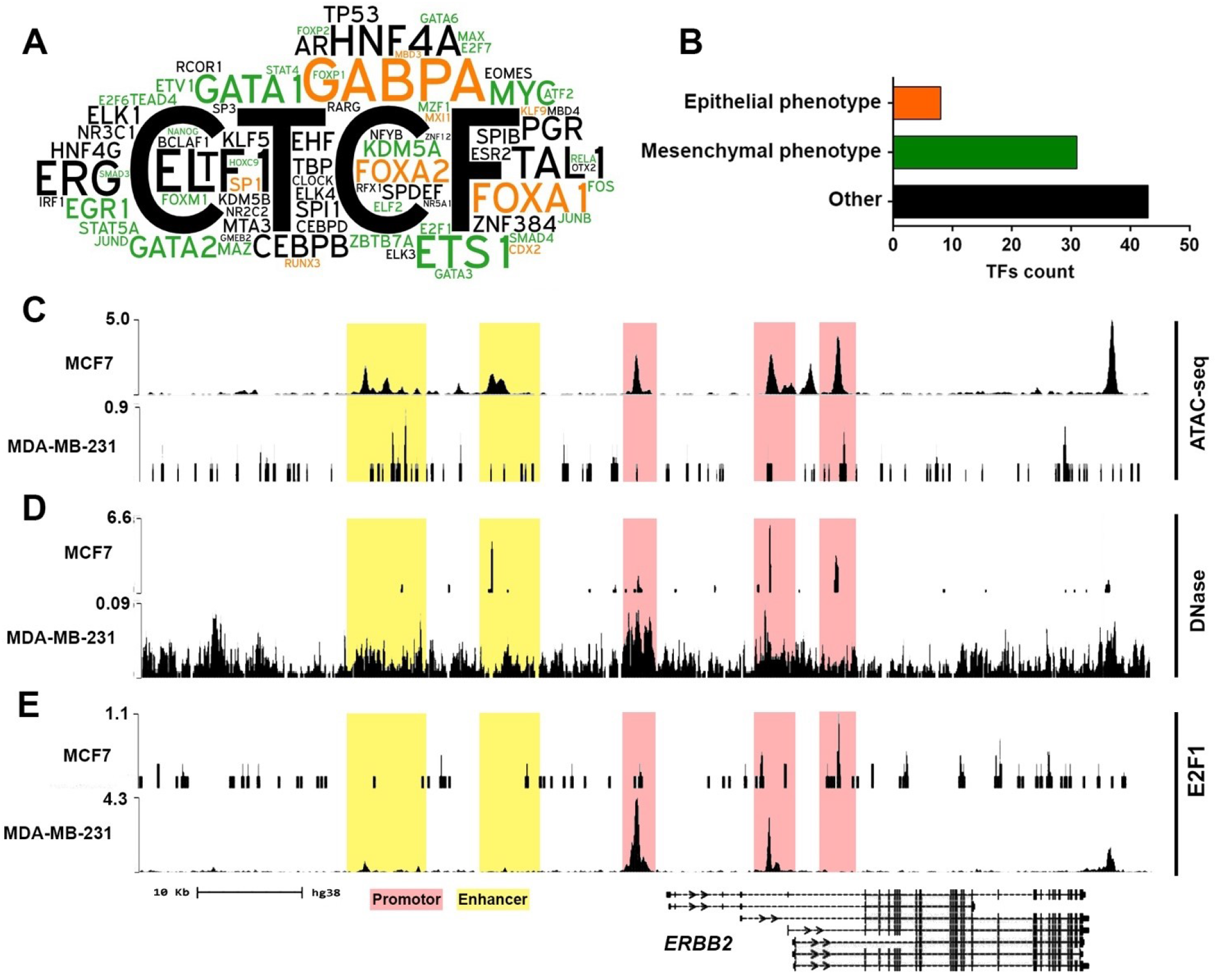
*ERBB2* gene binding transcription factors identified by ChIP-seq. (**A**) Word cloud diagram of transcription factors (TFs) detected bound to *ERBB2* gene at 10 kb upstream and downstream of motif Y. The different number of binding sites for each TF at the query region is illustrated as a different word size. TFs promoting epithelial and mesenchymal phenotypes are illustrated in orange and green colors respectively. (**B**) The number of the identified TFs based on their function in EMT procedure. Data obtained from ChIPbase v2.0 database available at “http://rna.sysu.edu.cn/chipbase/index.php”. (**C-E**) ChIP-seq enrichment value peaks of ATAC-seq (**C**), DNase I hypersensitivity (**D**) and E2F1 (**E**) at *ERBB2* gene regulatory regions in MCF7 and MDA-MB-231 cell lines. The normalized ChIP-seq values were obtained from Cistrome Data Browser [26] and the enrichment peaks were visualized by using WashU Epigenome Browser [27]. Genomic coordinate: chr17:39,643,771-39,735,523 (GRCh38/hg38 assembly).

We then investigated chromatin accessibility/activity of *ERBB2* regulatory elements (promoter and enhancer) by analyzing ATAC-seq and DNase hypersensitivity data of MCF7 and MDA-MB-231 cell lines to explore the mode of correlation of *ERBB2* expression with accessibility/activity of *ERBB2* regulatory element chromatin. The result showed higher ATAC-seq (Figure 5C) and DNase I hypersensitivity (Figure 5D) signals at *ERBB2* promoters and enhancer chromatin of MCF7 cell lines compared to MDA-MB-231 cell line. These results show that *ERBB2* promoter and enhancer chromatin of cells with higher *ERBB2* expression level is more accessible/active than that of cells with a lower level of *ERBB2* expression.

To examine whether higher *ERBB2* expression is also correlated with higher enrichment of mesenchymal phenotype transcription factor E2F1 at the regulatory regions of *ERBB* gene. Results revealed higher enrichment of E2F1 at promoter chromatin of *ERBB2* gene in the MA-MB-231 cell line was higher than that in MCF7 cell line (Figure 5E). These results demonstrate that lower *ERBB2* expression is correlated with higher enrichment of mesenchymal phenotype transcription factor at the regulatory elements of *ERBB2* gene.

### *ERBB2* chromatin histone marks in epithelial-like and mesenchymal-like breast cancer cells

To investigate whether different *ERBB2* expression in epithelial-like and mesenchymal-like cells is due to different *ERBB2* chromatin architecture, we studied the *ERBB2* chromatin dynamics in HER2-high (AU565, BT474, HCC-1954, MDA-MB-361, SKBR3) and HER2-low (MCF7, MDA-MB-231, MDA-MB-468) breast cancer cell lines. To this end, we analyzed the enrichment of open/active gene body chromatin histone marks (H2BK120ub, H3K39me3, H3K79me2), open/active promoter chromatin histone marks (H3K4me1, H3K4me3), open/active enhancer chromatin histone marks (H3K9ac, H3K27ac, H4K8ac), as well as closed/inactive promoter and enhancer chromatin histone marks (H3K9me, H3K27me3) at *ERBB2* gene chromatin in the cells. The normalized ChIP-seq values were obtained from Cistrome Data Browser [26] and the enrichment peaks were visualized by using WashU Epigenome Browser [27]. The cell lines were selected according to *ERBB2* expression levels shown in Figure 3A. The mRNA expression levels of the *ERBB2, MUC1, VIM,* and *GAPDH* genes are also shown in Figures 6A-D.

**Figure 6.**
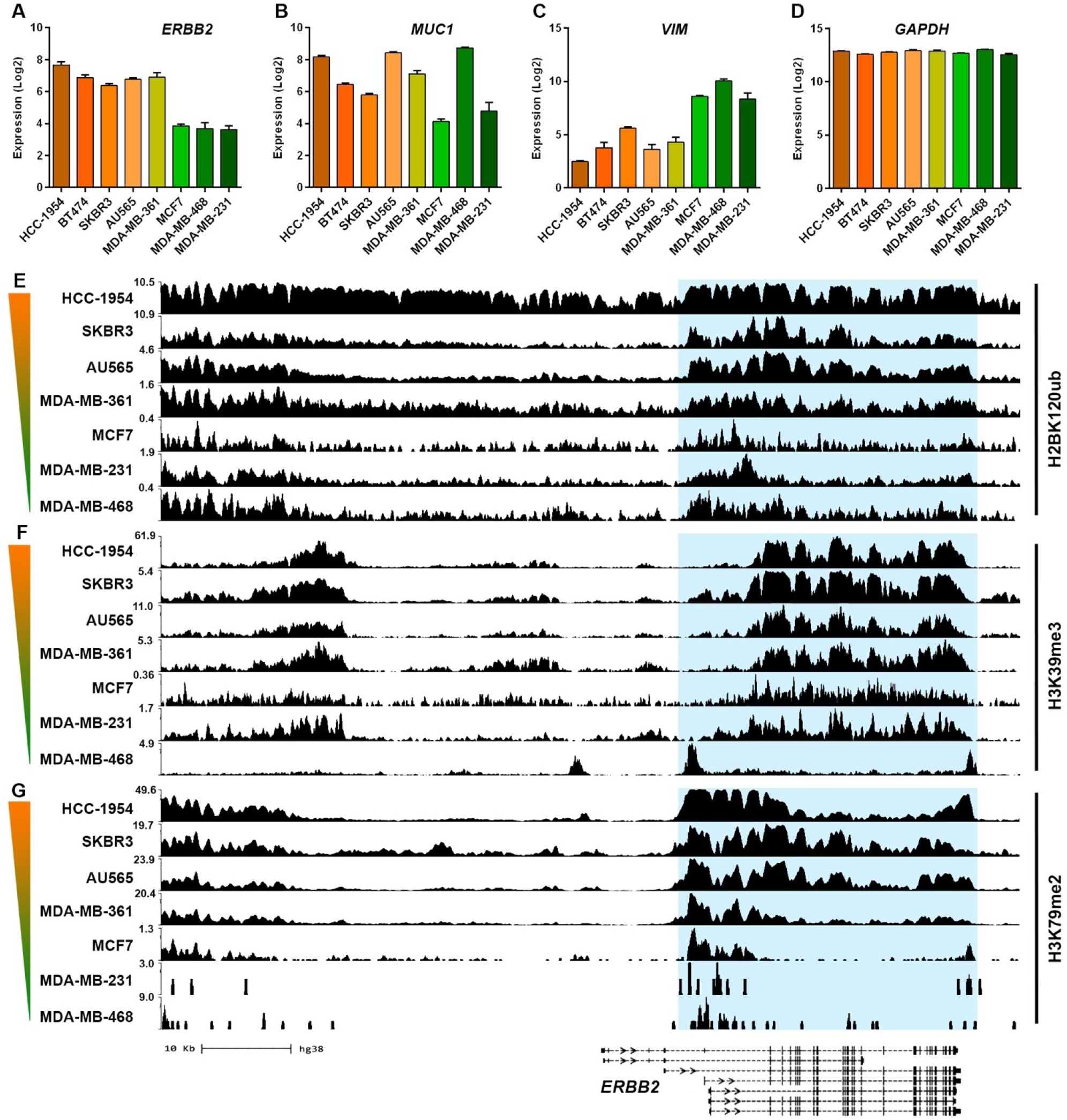
mRNA expression and open/active gene body chromatin histone marks of *ERBB2* gene in breast cancer cell lines. (**A-D**) mRNA expression levels of *ERBB2* (**A**), epithelial marker *MUC1* (**B**), mesenchymal marker *(VIM)* and GAPDH in HER2-high (HCC-1954, BT474, SKBR3, AU464, MDA-MB-361) and HER2-low (MCF7, MDA-MB-231, MDA-MB-468) breast cancer cell lines. Data obtained from GEO database with accession numbers GSE50811 [23] and GSE66071 [24]). (**E-G**) ChIP-seq enrichment of open/active gene body chromatin histone modification marks H2BK120ub (**E**), H3K39me3 (**F**), and H3K79me2 (**G**) at the chromatin of *ERBB2* gene and upstream region in the cell lines. Color gradient bars indicate the HER2 expression level. Genomic coordinate: chr17:39643771-39735523 (GRCh38/hg38 assembly).

As result, AU565, BT474, HCC-1954, MDA-MB-361 and SKBR3 cell lines showed higher *MUC1* mRNA expression compared to. In contrast, MCF7 and MDA-MB-231 but not MDA-MB-468 cell lines showed higher *VIM* expression compared to AU565, BT474, HCC-1954, MDA-MB-361 and SKBR3 cell lines (Figures 6A-D). These results further approve that *ERBB2* expression is correlated positively with epithelial phenotype and negatively with mesenchymal phenotype suggesting that mesenchymal-like breast cancer cells show low *ERBB2* expression.

ChIP-seq data of histone marks showed higher enrichment of H2BK120ub (Figure 6E), H3K39me5 (Figure 6F) and H3K79me3 (Figure 6G) at *ERBB2* gene body in the HER2-high cell lines compared to that in the HER2*-*low cell lines. Results also showed that enrichment of open/active promoter chromatin marks H3K4me2 (Figure 7A) and H3K4me3 (Figure 7B) at promoter chromatin of *ERBB2* gene in EHR2-high cell lines were significantly higher than that in HER2-low cell lines. The HER2-high cell lines showed relatively higher enrichment levels of open/active enhancer chromatin histone marks H3K9ac (Figure 8A), H3K27ac (Figure 8B) and H4K8ac (Figure 8C) at enhancer chromatin of *ERBB2* gene when compared with the HER2-low cell lines. In addition, enrichment levels of closed/inactive promoter and enhancer chromatin histone marks H3K9me (Figure 9A) and H3K27me3 (Figure 9B) at *ERBB2* gene were relatively low in HER2-high as well as in HER2-low cell lines.

**Figure 7.**
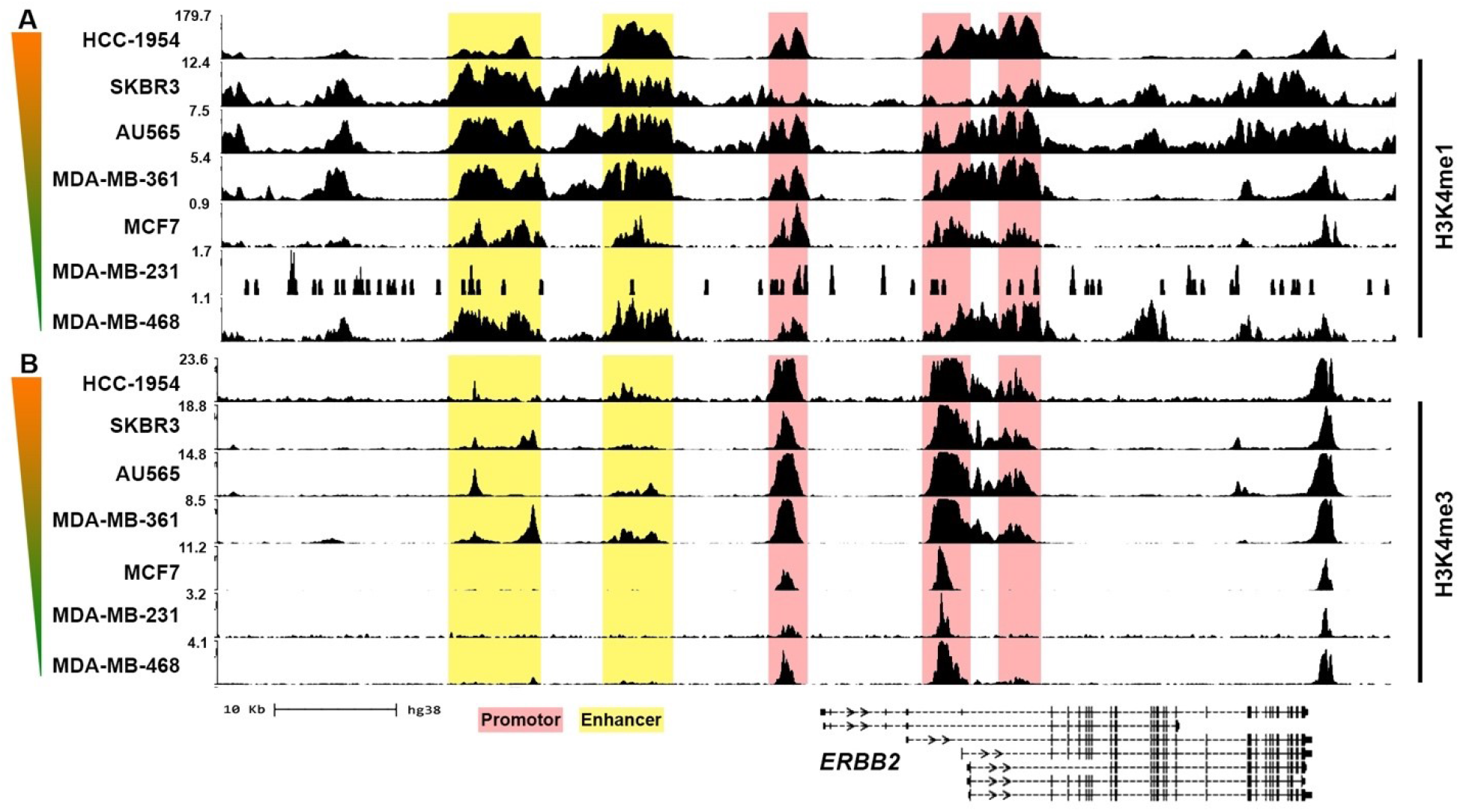
Open/active promoter histone marks of *ERBB2* gene in breast cancer cell lines. ChIP-seq enrichment of (**A**) H3K4me1 and (**B**) H3K4me3 at the chromatin of *ERBB2* gene and upstream region in HER2-high (HCC-1954, BT474, SKBR3, AU464, MDA-MB-361) and HER2-low (MCF7, MDA-MB-231, MDA-MB-468) breast cancer cell lines. Color gradient bars indicate the HER2 expression level. Genomic coordinate: chr17:39,643,771-39,735,523 (GRCh38/hg38 assembly).

**Figure 8.**
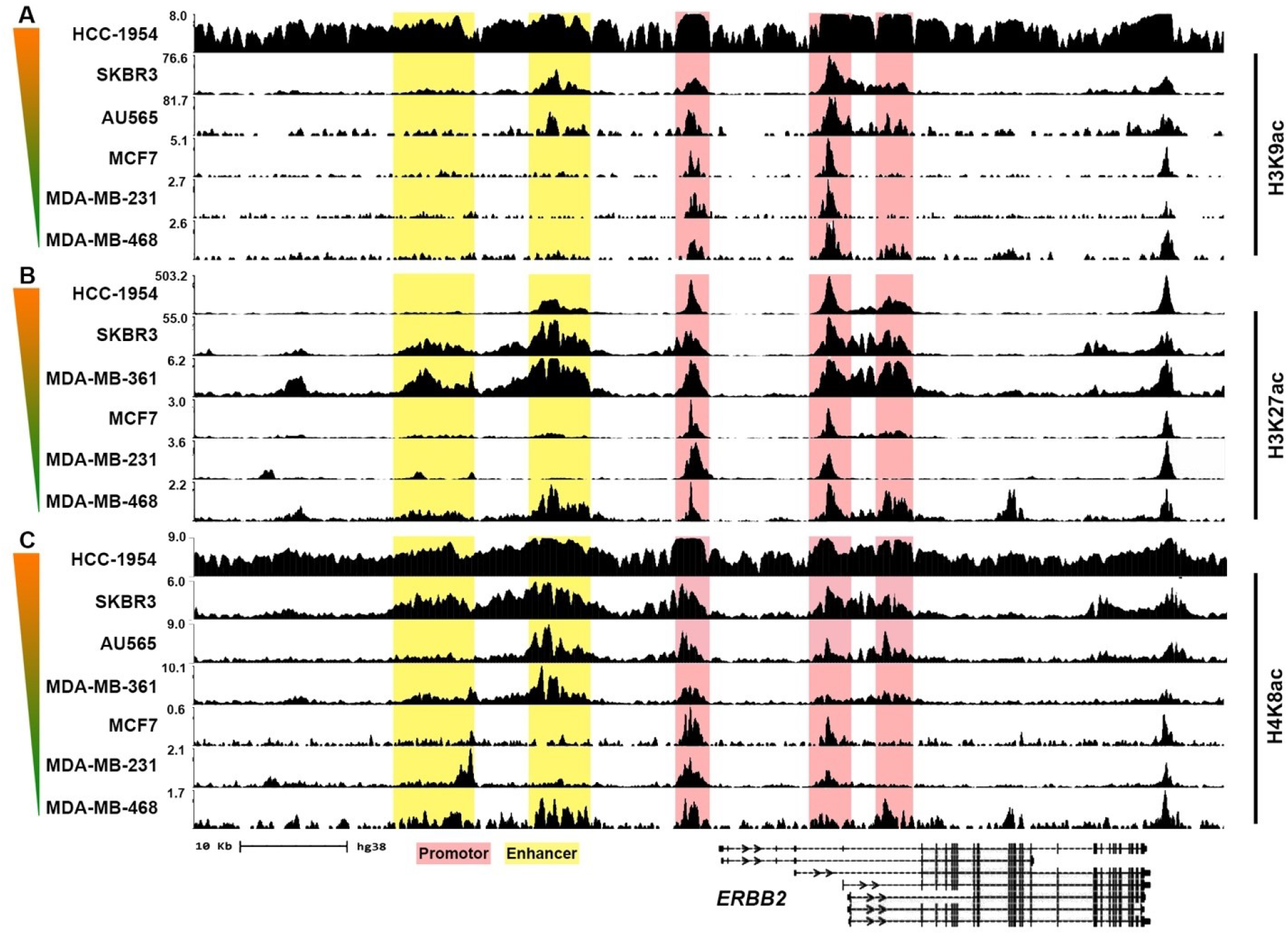
Open/active enhancer histone marks of *ERBB2* gene in breast cancer cell lines. ChIP-seq enrichment of (**A**) H3K9ac, (**B**) H3K27ac and (**C**) H4K8ac at the chromatin of *ERBB2* gene and upstream region in HER2-high (HCC-1954, BT474, SKBR3, AU464, MDA-MB-361) and HER2-low (MCF7, MDA-MB-231, MDA-MB-468) breast cancer cell lines. Color gradient bars indicate the HER2 expression level. Genomic coordinate: chr17:39,643,771-39,735,523 (GRCh38/hg38 assembly).

**Figure 9.**
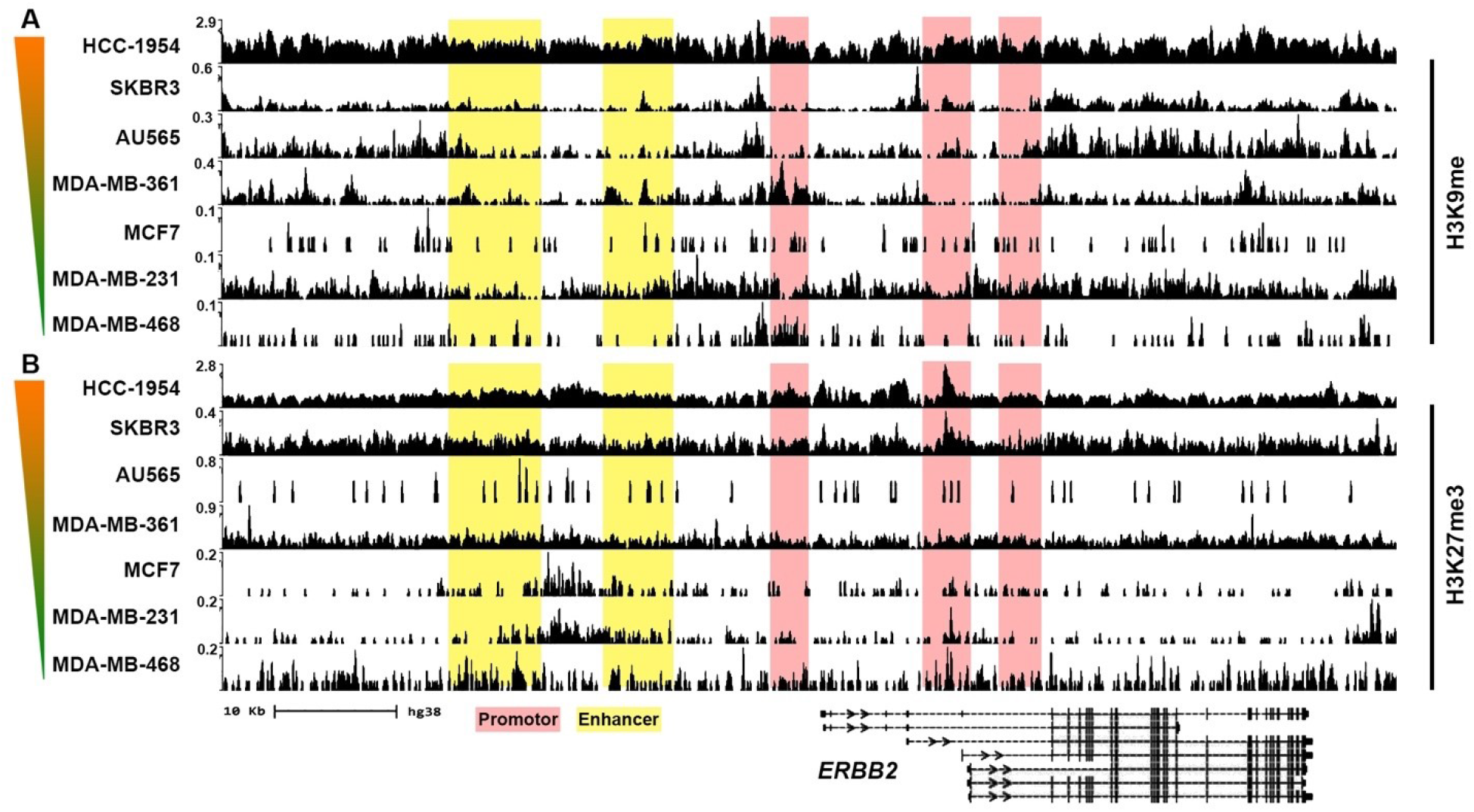
Closed/inactive promoter and enhancer histone marks of *ERBB2* gene in breast cancer cell lines. ChIP-seq enrichment of (**A**) H3K9me and (**B**) H3K27me3 at the chromatin of *ERBB2* gene and upstream region in HER2-high (HCC-1954, BT474, SKBR3, AU464, MDA-MB-361) and HER2-low (MCF7, MDA-MB-231, MDA-MB-468) breast cancer cell lines. Color gradient bars indicate the HER2 expression level. Genomic coordinate: chr17:39,643,771-39,735,523 (GRCh38/hg38 assembly).

Taken together, these results suggest that (1) Chromatin activity of the *ERBB2* gene governs *ERBB2* gene expression in breast cancer. *ERBB2* chromatin activity (2) Epithelial-like breast cancer cells express higher levels of *ERBB2* in comparison with mesenchymal-like breast cancer cells due to open chromatin of *ERBB2* promoter and enhancer regions in the epithelial-like breast cancers. *ERBB2* chromatin activity is correlated with the enrichment of epithelial phenotype transcription factors and open/active chromatin histone modifications. (3) Whereas mesenchymal-like cells show negligible *ERBB2* expression. This is because of closed/inactive *ERBB2* promoter and enhancer chromatin that is correlated with the absence of epithelial phenotype transcription factors as well with higher enrichment of mesenchymal phenotype transcription factors at their *ERBB2* chromatin. (4) Closed/inactive status of *ERBB2* chromatin in mesenchymal-like cells is not due to inactivator histone modifications, but maybe because of the absence of activator histone modifications at *ERBB2* chromatin.

### Chromatin-chromatin interaction of *ERBB2* in HER2-high and HER2-low breast cancer cell lines

We hypothesized that the interaction of *ERBB2* promoter with canonical and non-canonical enhancers is lower in HER2-low mesenchymal-like cells. To examine this, we performed a 4D genome organization analysis of *ERBB2* gene by analyzing interactions between *ERBB2* chromatin and upstream and downstream chromatin regions in HCC-1954 and MCF7 cell lines. To this objective, we analyzed experimental IM-PET (Integrated Methods for Predicting Enhancer Targets) and ChIA-PET (Chromatin Interaction Analysis by Paired-End Tag Sequencing) data from the cell lines by using 4Dgenome database [28]. Results showed the interaction of *ERBB2* promoter with 240 target enhancer regions in HCC-1954 cell line (Figures 10A and B). Of 240 target enhancers, 134 were at upstream and 106 were at downstream of *ERBB2* promoter. The chromatin loop size of 106 interactions was found smaller than 50 kb and 18 interactions formed a chromatin loop larger than 500 kb (Figures 10A and B). While MCF7 cell line showed the interaction of *ERBB2* promoter with 11 target enhancers which all were at upstream region of *ERBB2* promoter. Of 11 interactions, 10 had a chromatin loop size of fewer than 50 kb, and 1 interaction had a loop size of approximately 244 kb (Figures 10C and D).

**Figure 10.**
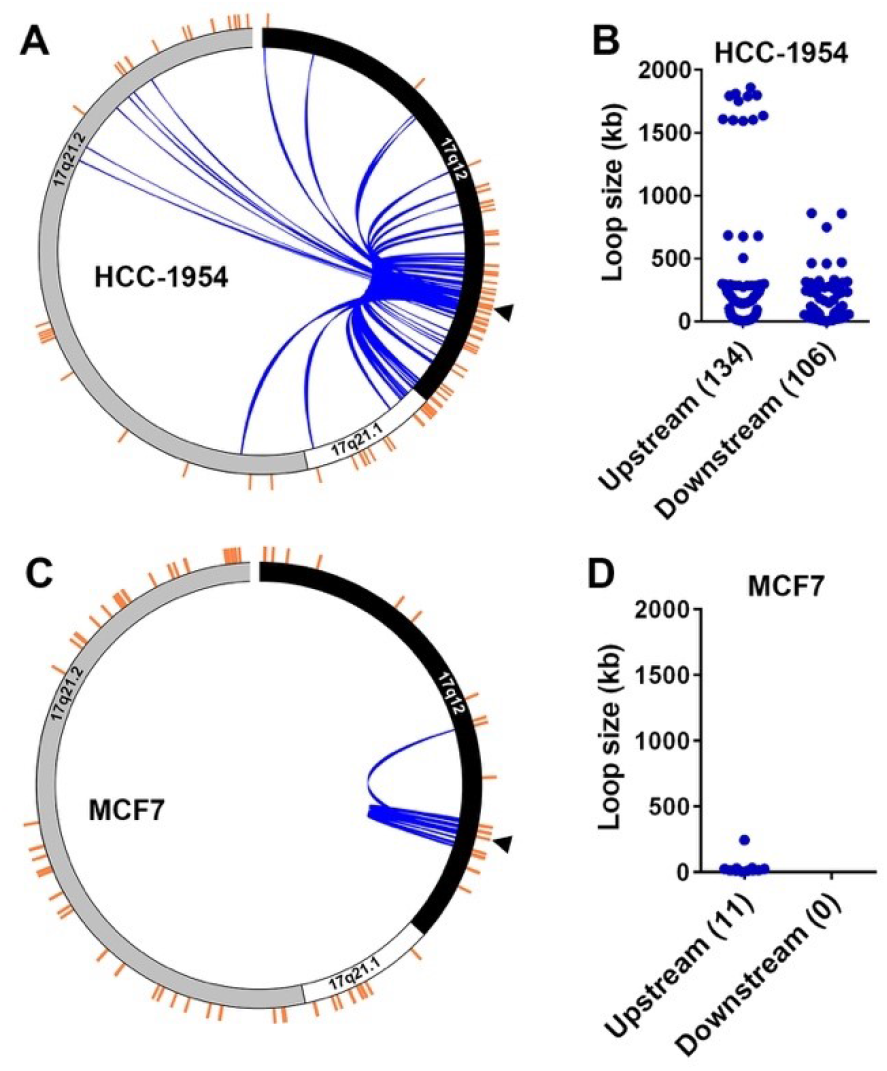
3D genome organization of *ERBB2* gene in breast cancer cell lines. (**A**) Circle plot of IM-PET promoter-enhancer interaction of *ERBB2* gene in HCC-1954 cell lines. (**B**) Scatter plot illustrating chromatin loop size of the *ERBB2* promoter-enhancer interactions divided by location of target enhancers (upstream or downstream from *ERBB2* TSS). (**C**) IM-PET promoter-enhancer interaction of *ERBB2* gene in MCF7 cell lines. (**D**) Scatter plot illustrating chromatin loop size of *ERBB2* promoter-enhancer interactions divided by location of target enhancers (upstream or downstream from *ERBB2* TSS). *ERBB2* TSS is indicated by arrowhead. Data was obtained from 4Dgenome database [28].

We also analyzed ChIP-seq H3K27ac enrichment profile of the *ERBB2* interaction sites in HCC-1954 and MCF7 in order to examine whether *ERBB2* promoter interaction with the target enhancer depends on chromatin activity of *ERBB2* promoter or target enhancer. The result showed higher H3K27ac enrichment at *ERBB2* chromatin in HCC-18954 cells in comparison with MCF7 cell lines. Whereas, MCF7 cell line showed higher H3K27ac enrichment at non-*ERBB2* chromatin than that in HCC-1954 cell lines. These results show the association of number and loop size of *ERBB2* chromatin interaction with the H3K27ac signature of ERBB2 chromatin but not non-ERBB2 chromatin. This suggests that ERBB2 promoter interaction with enhancers depends on the accessibility/activity of *ERBB2* promoter chromatin but not target enhancer chromatin. Overall, these results show that the number and size of promoter-enhancer interactions of *ERBB2* gene in HER2-high epithelial-like breast cancer cells are higher than that in HER2-low mesenchymal-like breast cancer cells. This is due to the accessibility/activity of *ERBB2* chromatin in HER2-high epithelial-like cells, and inaccessibility/inactivity of *ERBB2* chromatin in HER2-low mesenchymal-like cells.

Taken together, these results indicate higher 3D chromatin interaction of *ERBB2* gene in the HER2-high breast cancer cells compared to the HER2-low breast cancer cells. HER2-high epithelial-like breast cancer cells show a higher number of chromatin interactions between *ERBB2* promoter and target enhancers. Whereas the number of *ERBB2* promoter-enhancer interactions in HER2-low mesenchymal cells is lower. In addition, accessibility/activity of ERBB2 chromatin was the main factor in the formation of ERBB2 chromatin interaction with distance enhancer regions. This indicates the epigenetic role of breast cancer EMT regulators in the chromatin dynamics of *ERBB2* that controls HER2 expression in breast cancer cells.

### Downregulated *ERBB2* expression and upregulated EMT in lapatinib resistant cells

To investigate the role of EMT in development of resistance of HER2-positive breast cancer cells to anti-HER2 drugs, we studied the mRNA expression of *ERBB2*, epithelial phenotype markers *CDH1, ALCAM, FOXA1, NECTIN2* and *OCLN,* mesenchymal phenotype markers *CDH2, FN1, FOXC1, SNAI2* and *VIM,* as well as matrix metalloproteinase *MMP1, MMP2, MMP3, MMP9, MMP10* and *MMP28* in lapatinib sensitive and acquired lapatinib resistant BT474 cells. The array expression profiling data obtained from GEO database (Series GSE16179 [29]).

As result, lapatinib resistant cells showed lower expression levels of *ERBB2* (Figure 11A) and the epithelial marker genes (Figure 11B) and higher expression levels of the mesenchymal marker genes (Figure 11C) and MMPs (Figure 11D) when compared with the lapatinib sensitive cells. These results indicate that lapatinib sensitive cells are HER2-high epithelial-like cells with higher *ERBB2* gene expression, while lapatinib resistance cells are mesenchymal-like cells with lower HER2 expression compared to the lapatinib sensitive cells. These results suggest that EMT induces resistance to lapatinib via downregulating HER2 expression.

**Figure 11.**
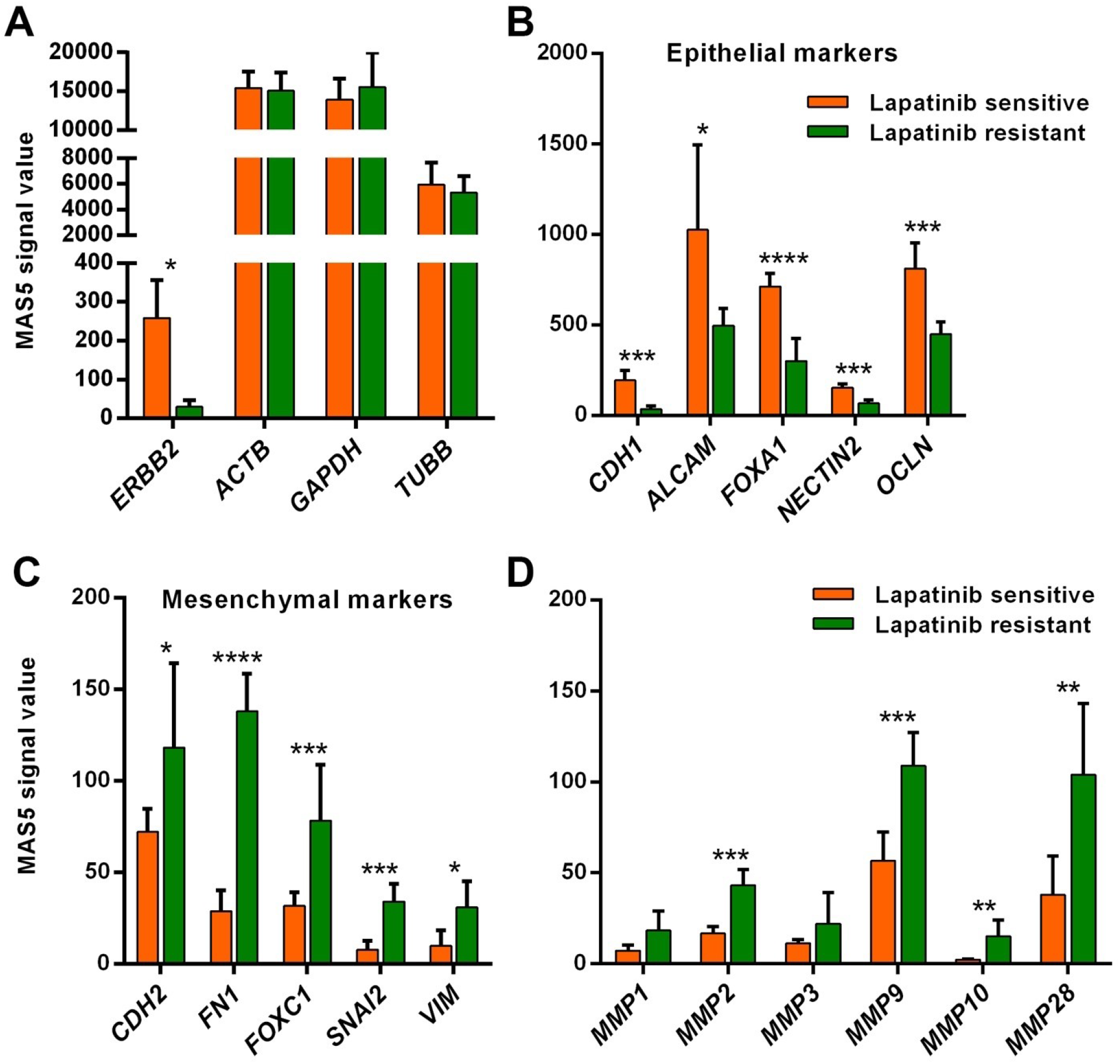
Downregulated *ERBB2* expression and upregulated EMT in lapatinib resistance. mRNA expression levels of (**A**) *ERBB2* and housekeeping genes, (**B**) epithelial phenotype marker genes, (**C**) mesenchymal phenotype marker genes and (**D**) matrix metalloproteinases in lapatinib sensitive and resistant BT474 cells. Data obtained from GEO database (Series GSE16179 [29]).

### EMT reduces HER2 expression and decreases trastuzumab binding to HER2-positive breast cancer cells

To examine whether induction of EMT of HER2-positive epithelial cell reduces HER2 expression, we analyzed mRNA expression of *ERBB2*, epithelial phenotype markers *CDH1, EPCAM, MUC1* and *OCLN,* mesenchymal phenotype markers *CDH2, FN1, SNAI2* and *VIM,* matrix metalloproteinase *MMP1, MMP2, MMP3, MMP9, MMP10* and *MMP28,* as well as *ADAM10, ADAM17* and *ADAM19* in A549 cell line (HER2-high human lung cancer epithelial cell line) subjected to EMT induction by treatment with 5 ng/ml TGF-β for 0, 0.5, 1, 2, 4, 8, 16, 24, and 72 hours. The array expression profiling data obtained from GEO database (Series GSE17708 [30]). Results showed significantly decreased mRNA expression of the epithelial marker genes (Figure 12) and increased mRNA expression of the mesenchymal marker genes (Figure 12), MMPs (Figure 12) as well as ADAMs (Figure 12), indicating EMT induction in the cells. Downregulated epithelial marker genes and upregulated mesenchymal marker genes were correlated with significant decline of *ERBB2* expression (*P* < 0.001) at 72 hours after TGF-β treatment started (Figure 12).

**Figure 12.**
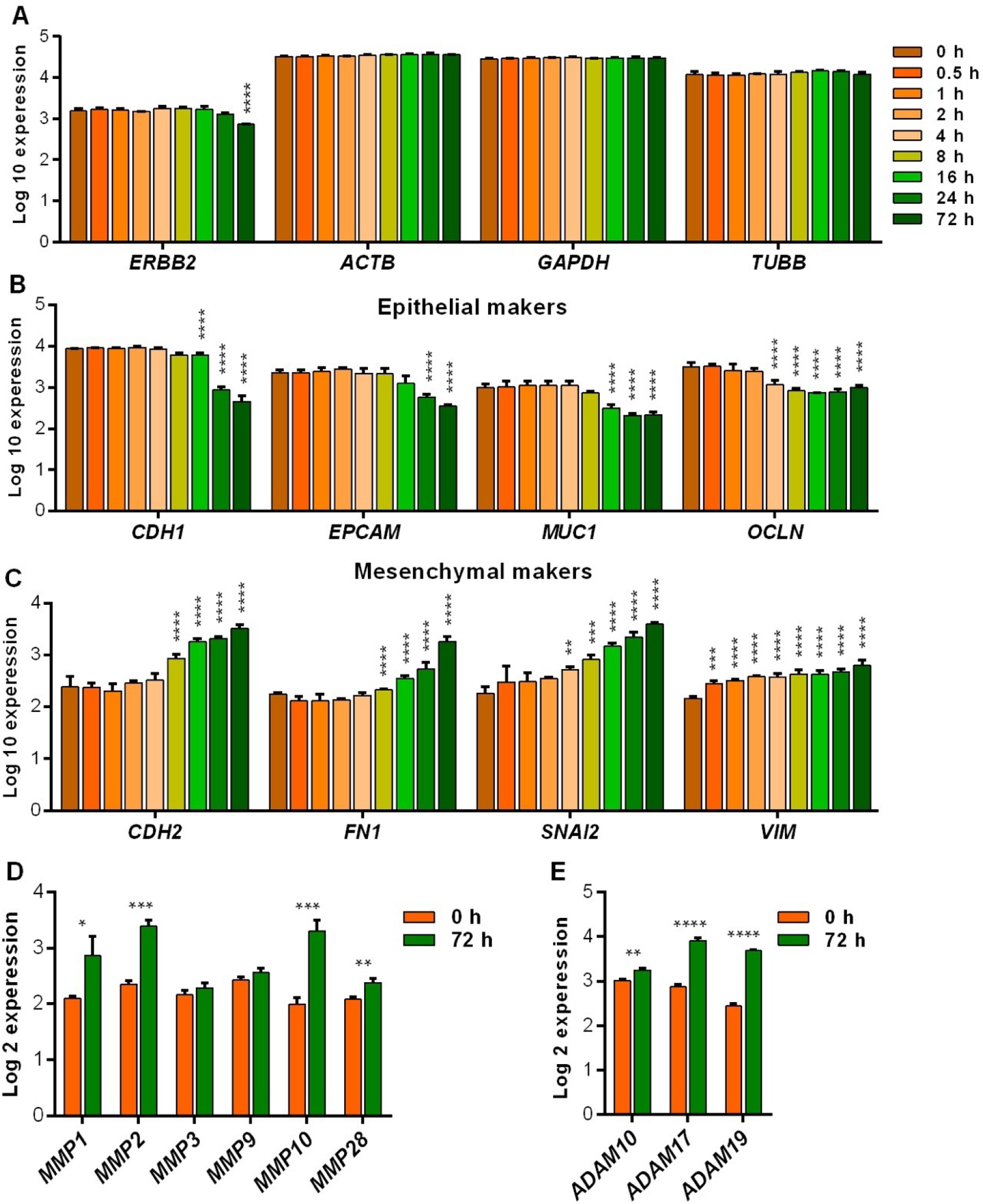
EMT reduces *ERBB2* expression. mRNA expression levels of (**A**) *ERBB2* and housekeeping genes, (**B**) epithelial phenotype marker genes, (**C**) mesenchymal phenotype marker genes and (**D**) MMPs and (**E**) ADAMs in TGF-β-mediated EMT-induced A549 cells. Data obtained from GEO database (Series GSE17708 [30]).

To investigate whether EMT reduces trastuzumab binding to HER2 we induced EMT in BT474 by treatment with EMT inducing media supplements cocktail (containing Wnt-5a, TGF-β1, antihuman E-Cadherin antibody, anti-human sFRP1 antibody, and anti-human Dkk1 antibody) for 15 days. Induction of EMT was confirmed by monitoring cell morphology (Figure 13) and immunofluorescence staining of Vimentin (Figure 13). After the majority of cells gained mesenchymal phenotype, the cells were treated with 10 μg/ml trastuzumab for 1 hour and then trastuzumab was stained by immunofluorescence staining. Results showed lower HER2 expression as well as lower binding of trastuzumab to HER2 in the cells underwent EMT compared to control cells. These results confirm that EMT downregulates HER2 expression that causes decreased rate of trastuzumab binding to HER2-positive breast cancer cells.

**Figure 13.**
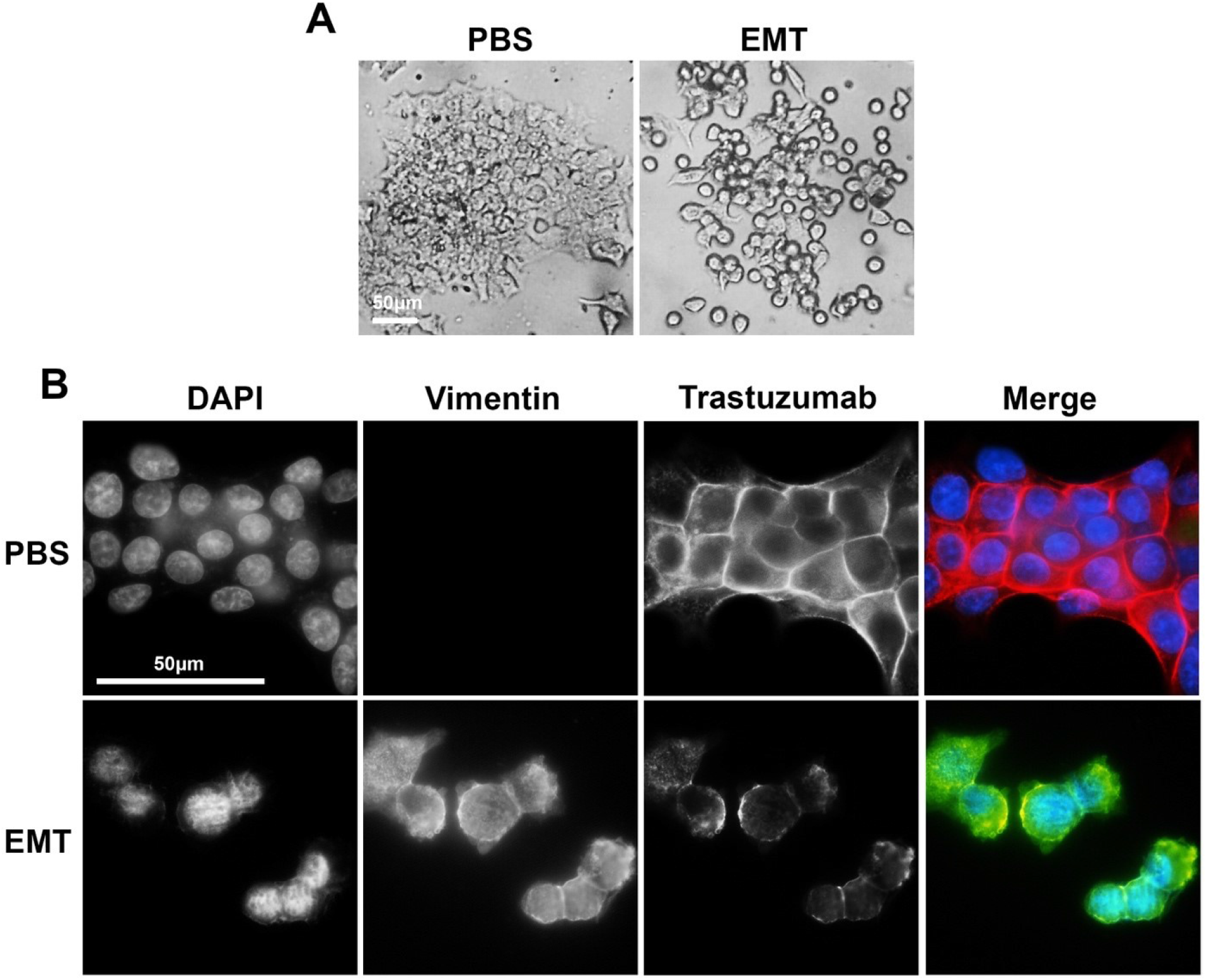
EMT decreases trastuzumab binding to HER2. (**A**) Epithelial morphology of PBS-treated BT474 cells and mesenchymal morphology of EMT-induced BT474 cells. (**B**) Immunofluorescence staining of Vimentin and trastuzumab in PBS-treated and EMT-induced BT474 cells treated with trastuzumab (10 μg/ml) for 1 hour.

## DISCUSSION

HER2 is an important target for the treatment of HER2-positive breast cancers. Several HER2-targeting agents such as trastuzumab and lapatinib have been approved by the FDA to treat HER2 positive breast cancer [12]. However, the resistance to these HER2 targeting agents has become a huge obstacle for the treatment of HER2-positive breast cancer patients. It is not clear why many HER2-positive tumors develop resistance to anti-HER2 neoadjuvant trastuzumab. Here, we studied the chromatin signature of *ERBB2* gene in epithelial-like HER2-high and mesenchymal-like HER2-low breast cancer cells and the role of EMT-mediated epigenetic regulation in *ERBB2* chromatin organization by analyzing genomics and epigenomics data from publicly available databases. We found that the expression of EMT marker and inducer is negatively correlated with HER2 expression, and positively correlated with trastuzumab and lapatinib resistance. HER2 expression in epithelial-like breast cancer cells is significantly higher than that in mesenchymal-like breast cancer cells. This is due to open/active chromatin of *ERBB2* gene in epithelial-like cells, as well as closed/inactive chromatin of *ERBB2* gene in mesenchymal-like breast cancer cells. This figure is also correlated with enrichment levels of EMT regulator transcription factor at the *cis*-regulatory regions of *ERBB2* gene. We also showed downregulated HER2 expression and upregulated EMT in BT474 cells resistance to lapatinib compared to lapatinib-sensitive cells. Induction of EMT in HER2-high epithelial-like breast cancer cells resulted in the downregulation of HER2 and decreased rate of trastuzumab binding to the cells. Our results suggest that EMT of HER2-positive breast cancer cells results in abrogation of HER2 expression by chromatin-base epigenetic silencing of *ERBB2* gene that leads to emergence of resistance to trastuzumab (Figure 14).

**Figure 14.**
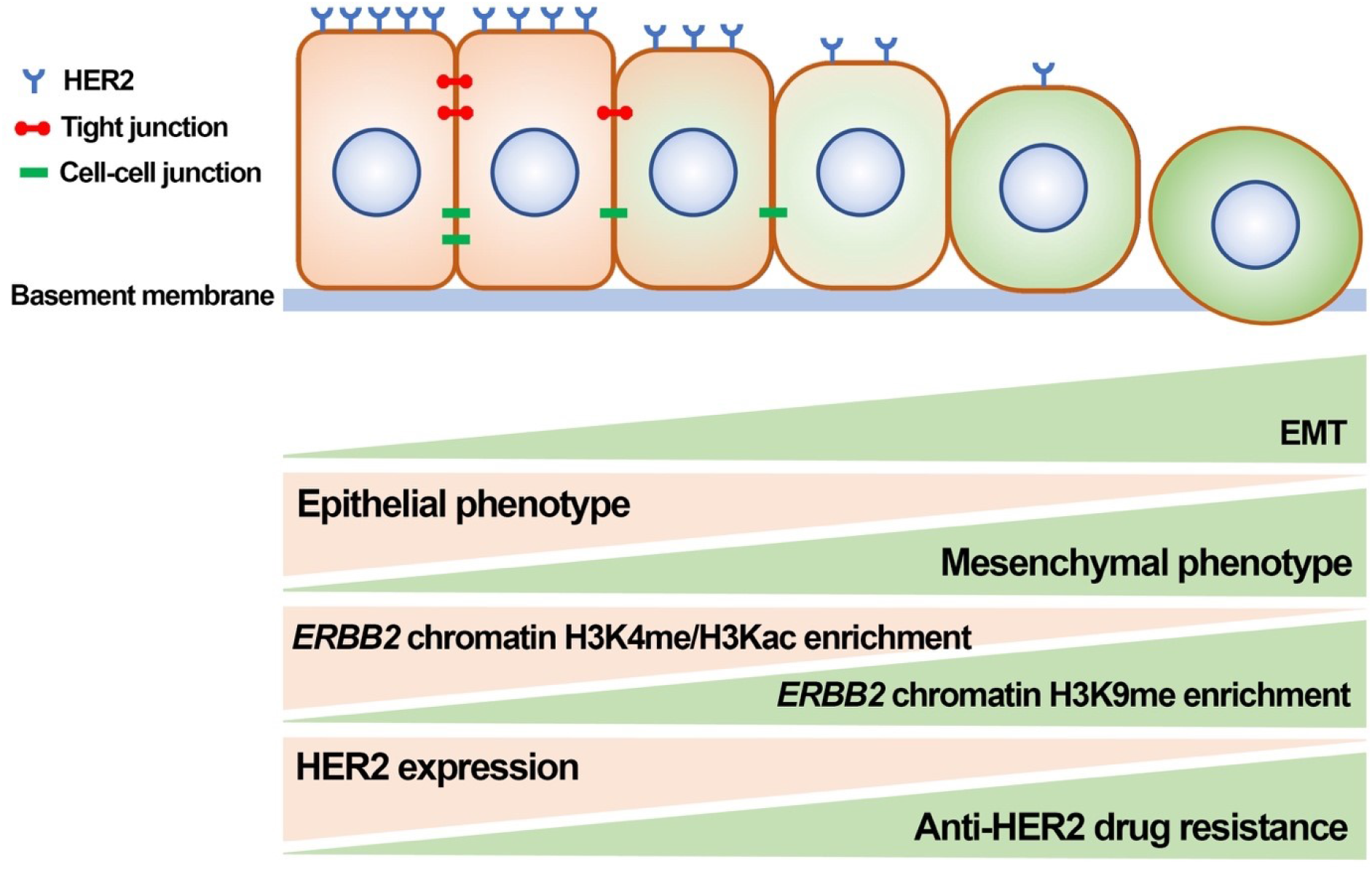
Schematic summary of findings. EMT of HER2-positive breast cancer cells increases trastuzumab resistance by chromatin-based epigenetic downregulation of HER2 expression. Increased EMT and mesenchymal phenotype is correlated with decreased expression of epithelial phenotype (including tight junctions and cell-cell junction proteins) and increased mesenchymal phenotype, decreased enrichment of open/active chromatin marks (H3K4me and H3Kac as examples), increased enrichment of closed/inactive chromatin marks (H3K9me as an example), decreased HER2 expression and increased trastuzumab resistance.

CD44+/CD24-cells are mesenchymal-like BCSCs that localized at the tumor invasive margins and are correspond to migration and metastasis, whereas ALDH+ cells are defined as epithelial-like BCSCs that are located in deeper sites of the tumors and exhibit more proliferative property [12,31]. CD44+/CD24- cells were isolated by FACS from non-tumorigenic human mammary epithelial cells that have undergone an induced EMT, exhibited many properties of BCSCs including mammosphere-formation ability [32]. On the other side, CD44+/CD24- BCSCs isolated from breast tumors expressed a low level of E-cadherin, but high levels of EMT markers including N-cadherin, Vimentin, Fibronectin, ZEB1/2, FOXC2, Snail, Slug, and Twist1/2 [32]. Clinical studies revealed that HER2-positive metastatic breast cancers were associated with EMT [33,34]. HER2 signaling in human breast epithelial cells results in declined expression of Vimentin, N-cadherin, and Integrin-α5, as well as the loss of E-cadherin and Desmoplakin. However, some recent study suggests that loss of E-cadherin is not essential for HER2-induced EMT [35,36]. It is shown recently that overexpression of HER2 in epithelial breast cancer cell line D492 induces EMT and maintains the mesenchymal phenotype in the absence of EGFR [37]. There are also many reports revealing crosstalk of HER2 receptor and its downstream pathways with the stemness signaling pathways to prone mammary epithelial cells towards EMT [12].

We previously reviewed the hypothesized that response to trastuzumab in luminal cells and resistance to trastuzumab in basal/mesenchymal cells may link to EMT [12]. Trastuzumab-resistant tumors are thought to be enriched for EMT features. Basal breast cancer cell line JIMT-1 which is HER2-positive, trastuzumab-refractory, ER-negative, and Vimentin-positive is a best cell model to explain the link between HER2, EMT and trastuzumab resistance. De novo resistance of JIMT-1 cells to trastuzumab can be explained by the emergence of trastuzumab-resistance BCSCs due to the dynamic interaction between HER2 and EMT [19,38–43]. The subpopulation of CD44+/CD24- BCSC in the early culture of the JIMT-1 cell line is reported around 10%. However, this rate was increased to 85% at the late-passages which was also associated with a dramatic decline in HER2 expression levels [19]. Treatment of CD44+/CD24- BCSCs derived from breast tumor tissues treated with formestane, an aromatase inhibitor, resulted in a 16% decrease (*P* < 0.01) in the cell proliferation in response to single-agent trastuzumab and 50% decrease (*P* < 0.001) in response to treatment with trastuzumab combined with formestane. The combined treatment also inhibits the expression of EGFR, HER2, aromatase and Cyclin D1 in CD44+/CD24- cells, which suggests that targeting HER2 by trastuzumab may inhibit the growth of CD44+/CD24- BCSCs through the inhibition of cell cycle progression [41]. Some other reports suggest that preferential killing of the putative CD44+/CD24- BCSCs might be sufficient to overcome primary resistance to trastuzumab. The CD44+/CD24- BCSCs derived from trastuzumab-refractory JIMT-1 cells were 10-fold more sensitive to cell growth inhibitory effects of metformin than the other cells [43]. JIMT-1 cell line is highly enriched with the mesenchymal phenotype CD44+/CD24- fraction in late passages [19,44,45]. Indeed, treatment of JIMT-1 tumors with trastuzumab failed to exhibit significant reductions in tumor volume but when trastuzumab was combined with metformin, the tumor size was significantly smaller than those of the groups treated with single-agent trastuzumab or metformin [43]. These results suggest that BCSCs inside JIMT-1 tumor escape from trastuzumab effects, that support the hypothesis that mesenchymal cell subpopulation is responsible for resistance of the tumor against trastuzumab.

It is further revealed that trastuzumab-resistant HER2-positive cells show spontaneous EMT and predominant exhibition of CD44+/CD24- phenotype [46]. There is a synchronous increase in CD44 and elements of Wnt/β-catenin signaling and a decrease in CD24 expression in mesenchymal colony clusters of SKBR3 cells. SKBR3 cell line is characterized as HER2-positive, trastuzumab-sensitive liminal cell line. The CD44+/CD24- mesenchymal colonies of SKBR3 also show significantly upregulated EMT markers including Vimentin, N-cadherin, Twist1 and Fibronectin. The colonies were highly resistant to trastuzumab and lapatinib while the luminal/epithelial SKBR3 cells remained trastuzumab-sensitive. Similar to previous reports, HER2 expression levels in mesenchymal colonies were negatively correlated with trastuzumab resistance [46]. Thus, lapatinib which is recently approved to treat trastuzumab-resistant breast cancers, may not be a useful therapeutic option in targeting CD44+/CD24- mesenchymal cell-rich tumors due to the negative regulation of HER2 during EMT. Furthermore, the expression of the EMT-driving transcription factors Slug, Twist1 and ZEB1 are higher in trastuzumab-refractory basal HER2-positive JIMT-1 cells than that in the trastuzumab-responsive SKBR3 cells. The knockdown of the transcription factors in parental JIMT-1 cells reduced the subpopulation of CD44+/CD24- BCSC by 5, 5 and 2-fold, respectively. Interestingly, depletion of the EMT-driving transcription factors increased the trastuzumab-refractory in JIMT-1 tumors due to sensitized CD44+/CD24- BCSCs inside the bulk JIMT-1 tumors [42]. HER2-positive basal epithelial BCSCs are susceptible to change in their expression signature during EMT. This phenomenon may explain the resistance of some HER2- positive breast tumors to HER2-targeting agent including trastuzumab. Further, CD44+/CD24- BCSC subpopulation of HER2-positive cell lines or tumors may escape from trastuzumab-mediated ADCC. BCSCs could survive the immunoselection process in breast cancer cells co-cultured with NK cells and trastuzumab. This resistance may be attributed to the reduced HER2 expression levels on their surface [20]. Overall, our results demonstrate that EMT of HER2-positive breast cancer cells causes the emergence of a mesenchymal-like cell subpopulation which are HER2-low/negative and highly resistant to trastuzumab and lapatinib. The significant depletion of HER2 in the mesenchymal-like cells inside the tumor can take place by chromatin-based epigenetic silencing of *ERBB2* gene during EMT. Thus, we suggest that EMT and mesenchymal-like cells can be major mechanism of and responsible for de novo resistance to HER2-targeting therapeutics including trastuzumab and lapatinib and serve as potential effective targets for therapy and to overcome drug resistance in breast cancers.

It has been demonstrated that HER2 itself induces EMT of breast cancer cells via upregulating stemness pathways. This suggests a negative feedback loop between HER2 and EMT that gives important clue about the trastuzumab-responsive HER2-positive breast cancer develops resistance to trastuzumab. Our results also suggest that *ERBB2* gene silencing by epigenetic regulation during EMT is an authentic mechanism of downregulated HER2 in the mesenchymal-like cells and the main mechanism of resistance of HER2-positive breast cancer cells to trastuzumab and lapatinib. As a future direction, we recommend investigating that mechanism of the negative feedback loop to understand how HER2 overexpression induces EMT, how EMT causes *ERBB2* gene silencing leading to emergence of tumor cells resistant to HER2-targeted therapies.

## METHODS

### Antibodies

Trastuzumab (Herceptin®) was purchased from Roche (Basel, Switzerland). Mouse monoclonal anti-human Vimentin (V9; cat# sc-6260) and FITC-conjugated rabbit anti-mouse (cat# sc-358916) antibodies was purchased from Santa Cruz Biotechnology (Dallas, TX, USA). Rhodamine (TRITC)- conjugated donkey anti-human IgG (Cat# 709-025-149) was from Jackson ImmunoResearch (West Grove, PA, USA).

### Cell culture

HER2-positive breast cancer cell line BT474 were purchased from American Type Culture Collection (ATCC; Manassas, VA, USA). The cells were cultured in Dulbecco’s modified Eagle’s medium (DMEM) medium supplemented with 10% fetal bovine serum (FBS) and antibiotics including penicillin (100 U/ml) and streptomycin (100 μg/ml) and were maintained at 5% CO2 atmosphere at 37°C. For treatment experiments, appropriate number of cells were seeded in DMEM medium containing 10% FBS (starvation medium) and were cultured for 24 hours. The cells then were starved overnight (16 hours) at DMEM containing 1% FBS before the treatments and then were cultured in starvation medium containing the test agent.

### EMT induction

Number of 2 × 10^4^ BT474 cells were seeded in each 24-well plate and cultured in DMEM medium containing 10% FBS and 1X StemXVivo EMT inducing supplement containing recombinant human Wnt-5a, recombinant human TGF-β1, anti-human E-cadherin, anti-human sFRP-1 and antihuman Dkk-1 antibodies (cat# CCM017; R&D Systems; Minneapolis, MN, USA) for 15 days. After treatment, successful EMT induction was studied by monitoring cell morphology and immunofluorescence staining of Vimentin.

### Immunofluorescence staining assay

The indirect double-immunofluorescence staining was done as described previously [47]. BT474 cells were cultured on 25 mm coverslip glasses in 24-well plate for 24 hours and then were treated with 10 μg/ml trastuzumab for an hour. Afterward, the coverslips were washed with ice-cold PBS and the cells were fixed by incubation in −20°C methanol for 5 minutes. The coverslips were then washed with TBS and blocked in coverslip blocking buffer (1% BSA solution in TBS) for 1 hour. After blocking, the coverslips were incubated in 2 μg/ml primary antibody (anti-human Vimentin) solution for 1 hour. The coverslips were washed and then incubated in 1 μg/ml FITC-conjugated and/or 1 μg/ml TRITC-conjugated secondary antibodies solutions for 1 hour in dark. Afterward, the coverslips were washed with TBS and were then incubated in 1 μg/ml DAPI solution for 5 minutes. The coverslips were mounted on microscope slides, sealed by nail polish and observed under a fluorescence microscope using FITC and TRITC channels.

### cBioPortal cancer genomics database

RNA-seq and expression Z-scores of 1,904 breast cancer tumors studied by METABRIC study were obtained from and analyzed using cBioPortal cancer genomics database [22,48] available at http://cbioportal.org/index.do.

### Gene Expression Omnibus (GEO)

All mRNA expression and methylation data from cell lines were obtained from GEO database available at https://www.ncbi.nlm.nih.gov/geo. GEO series and samples accession IDs of analyzed data are shown in Table 1.

**Table 1.**
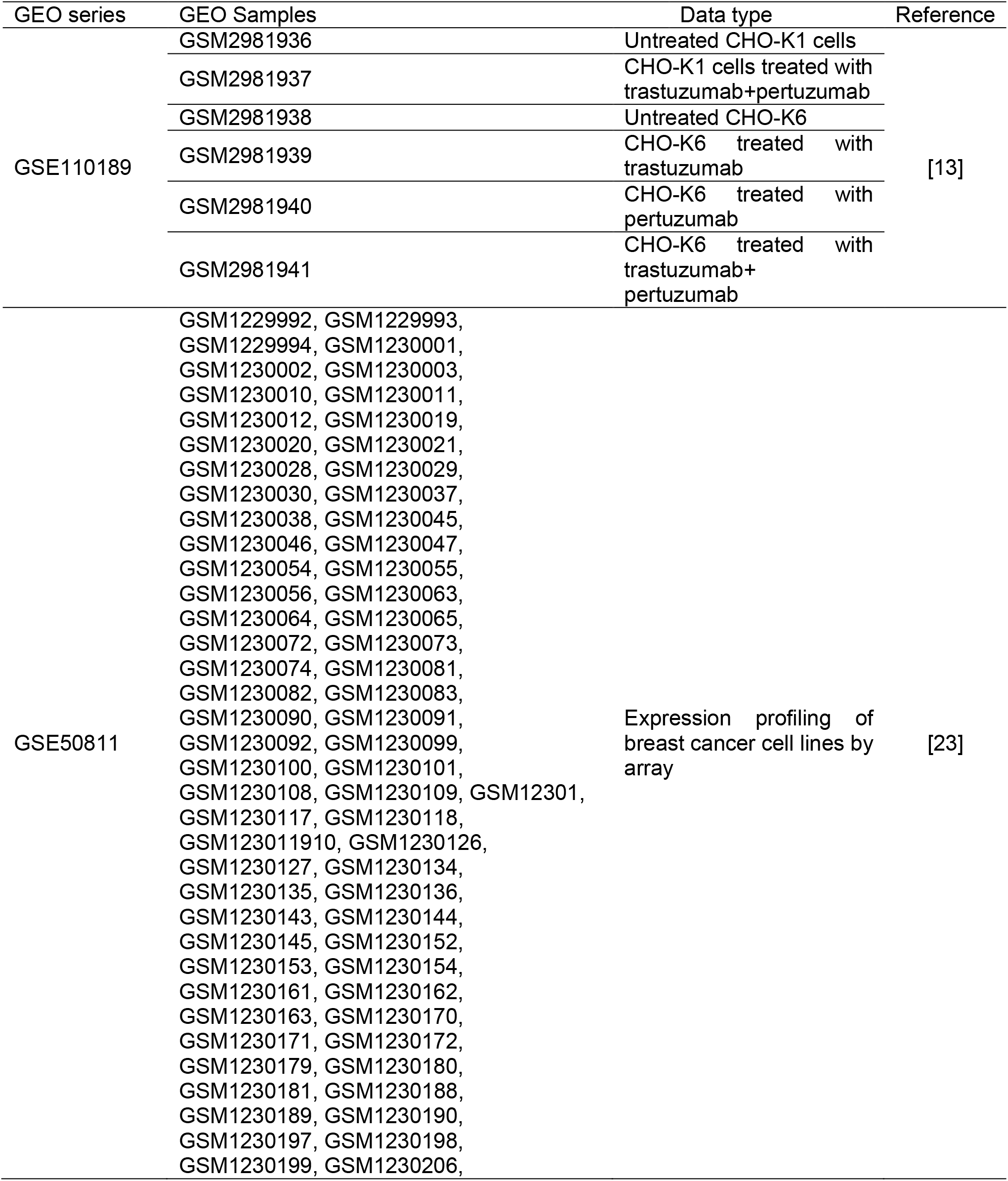
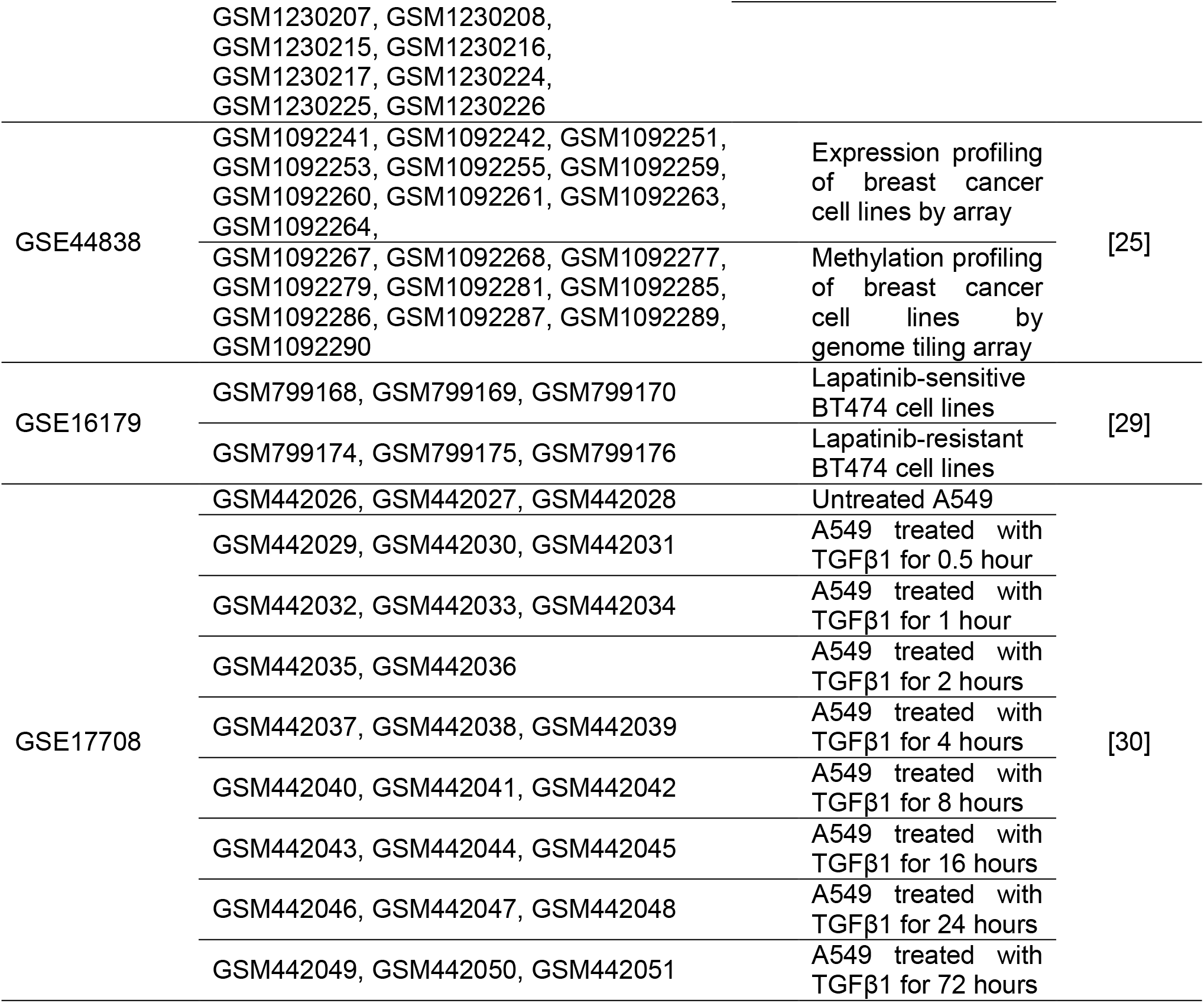
GEO series and samples accession IDs of expression and methylation arrays data analyzed in this thesis.

### ChIP-seq chromatin enrichment

ChIP-seq data were obtained from Cistrome Data Browser [26] available at http://cistrome.org/db/# and GEO database. ChIP-seq data were visualized by using WashU Epigenome Browser [27] available at “https://epigenomegateway.wustl.edu”. Cistrome DB and GEO samples accession IDs of analyzed ChIP-seq data are shown in Table 2.

**Table 2.** Cistrome DB and GEO accession IDs of ChIP-seq data analyzed in this thesis.

| Factor | X-seq | Cell line | Cistrome DB ID | GEO sample ID | Reference |
| --- | --- | --- | --- | --- | --- |
| Tn5 | ATAC-seq | MCF7 | 79676 | GSM2714245 | [49] |
| Tn5 | ATAC-seq | MDA-MB-231 | 65718 | GSM2439559 | [50] |
| DNase | DNase-seq | MCF7 | 40995 | GSM1008581 | [51] |
| DNase | DNase-seq | MDA-MB-231 | 78267 | GSM2242137 | [52] |
| FOXA1 | ChIP-seq | MDA-MB-453 | 36842 | GSM1099031 | [53] |
| FOXA1 | ChIP-seq | MCF7 | 2320 | GSM659787 | [54] |
| E2F1 | ChIP-seq | MCF7 | 2281 | GSM699986 | [55] |
| E2F1 | ChIP-seq | MDA-MB-231 | 75043 | GSM2501567 | [52] |
| H2BK120ub | ChIP-seq | HCC-1954 | 85603 | GSM2258929 | [56] |
| H2BK120ub | ChIP-seq | SKBR3 | 82286 | GSM2258950 | [56] |
| H2BK120ub | ChIP-seq | AU565 | 85597 | GSM2258923 | [56] |
| H2BK120ub | ChIP-seq | MDA-MB-361 | 81479 | GSM2258935 | [56] |
| H2BK120ub | ChIP-seq | MCF7 | 86405 | GSM2258947 | [56] |
| H2BK120ub | ChIP-seq | MDA-MB-231 | 81472 | GSM2258932 | [56] |
| H2BK120ub | ChIP-seq | MDA-MB-468 | 86411 | GSM2258941 | [56] |
| H3K39me3 | ChIP-seq | HCC-1954 | 88403 | GSM2258835 | [56] |
| H3K39me3 | ChIP-seq | SKBR3 | 87169 | GSM2258798 | [56] |
| H3K39me3 | ChIP-seq | AU565 | 88152 | GSM2258816 | [56] |
| H3K39me3 | ChIP-seq | MDA-MB-361 | 81935 | GSM2258762 | [56] |
| H3K39me3 | ChIP-seq | MCF7 | 74303 | GSM2483406 | [56] |
| H3K39me3 | ChIP-seq | MDA-MB-231 | 2408 | GSM425485 | [57] |
| H3K39me3 | ChIP-seq | MDA-MB-468 | 82822 | GSM2258889 | [56] |
| K3K79me2 | ChIP-seq | HCC-1954 | 85976 | GSM2258841 | [56] |
| K3K79me2 | ChIP-seq | SKBR3 | 84023 | GSM2258805 | [56] |
| K3K79me2 | ChIP-seq | AU565 | 84313 | GSM2258823 | [56] |
| K3K79me2 | ChIP-seq | MDA-MB-361 | 81942 | GSM2258769 | [56] |
| K3K79me2 | ChIP-seq | MCF7 | 86291 | GSM2258733 | [56] |
| K3K79me2 | ChIP-seq | MDA-MB-231 | 88654 | GSM2258858 | [56] |
| K3K79me2 | ChIP-seq | MDA-MB-468 | 87026 | GSM2258894 | [56] |
| H3K4me1 | ChIP-seq | HCC-1954 | 88402 | GSM2258836 | [56] |
| H3K4me1 | ChIP-seq | SKBR3 | 84019 | GSM2258801 | [56] |
| H3K4me1 | ChIP-seq | AU565 | 88154 | GSM2258818 | [56] |
| H3K4me1 | ChIP-seq | MDA-MB-361 | 81941 | GSM2258764 | [56] |
| H3K4me1 | ChIP-seq | MCF7 | 82432 | GSM2258728 | [56] |
| H3K4me1 | ChIP-seq | MDA-MB-231 | 68383 | GSM2036932 | [56] |
| H3K4me1 | ChIP-seq | MDA-MB-468 | 87022 | GSM2258890 | [56] |
| H3K4me3 | ChIP-seq | HCC-1954 | 8399 | GSM721134 | [58] |
| H3K4me3 | ChIP-seq | SKBR3 | 82540 | GSM2258803 | [56] |
| H3K4me3 | ChIP-seq | AU565 | 84311 | GSM2258821 | [56] |
| H3K4me3 | ChIP-seq | MDA-MB-361 | 81939 | GSM2258766 | [56] |
| H3K4me3 | ChIP-seq | MCF7 | 86289 | GSM2258731 | [56] |
| H3K4me3 | ChIP-seq | MDA-MB-231 | 68122 | GSM1700393 | [59] |
| H3K4me3 | ChIP-seq | MDA-MB-468 | 54526 | GSM1429760 | [60] |
| H3K9ac | ChIP-seq | HCC-1954 | 84601 | GSM2258842 | [56] |
| H3K9ac | ChIP-seq | SKBR3 | 84020 | GSM2258806 | [56] |
| H3K9ac | ChIP-seq | AU565 | 84254 | GSM2258824 | [56] |
| H3K9ac | ChIP-seq | MCF7 | 86292 | GSM2258734 | [56] |
| H3K9ac | ChIP-seq | MDA-MB-231 | 58110 | GSM1619768 | [61] |
| H3K9ac | ChIP-seq | MDA-MB-468 | 87027 | GSM2258897 | [56] |
| H3K27ac | ChIP-seq | HCC-1954 | 88400 | GSM2258830 | [56] |
| H3K27ac | ChIP-seq | SKBR3 | 87173 | GSM2258794 | [56] |
| H3K27ac | ChIP-seq | MDA-MB-361 | 85478 | GSM2258758 | [56] |
| H3K27ac | ChIP-seq | MCF7 | 82438 | GSM2258722 | [56] |
| H3K27ac | ChIP-seq | MDA-MB-231 | 61914 | GSM1855992 | [62] |
| H3K27ac | ChIP-seq | MDA-MB-468 | 82828 | GSM2258884 | [56] |
| H4K8ac | ChIP-seq | HCC-1954 | 81475 | GSM2258931 | [56] |
| H4K8ac | ChIP-seq | SKBR3 | 82288 | GSM2258952 | [56] |
| H4K8ac | ChIP-seq | AU565 | 85599 | GSM2258925 | [56] |
| H4K8ac | ChIP-seq | MDA-MB-361 | 81477 | GSM2258937 | [56] |
| H4K8ac | ChIP-seq | MCF7 | 86413 | GSM2258949 | [56] |
| H4K8ac | ChIP-seq | MDA-MB-231 | 81478 | GSM2258934 | [56] |
| H4K8ac | ChIP-seq | MDA-MB-468 | 86409 | GSM2258943 | [56] |
| H3K9me3 | ChIP-seq | HCC-1954 | 84600 | GSM2258845 | [56] |
| H3K9me3 | ChIP-seq | SKBR3 | 84024 | GSM2258808 | [56] |
| H3K9me3 | ChIP-seq | AU565 | 84308 | GSM2258826 | [56] |
| H3K9me3 | ChIP-seq | MDA-MB-361 | 84978 | GSM2258772 | [56] |
| H3K9me3 | ChIP-seq | MCF7 | 86294 | GSM2258736 | [56] |
| H3K9me2 | ChIP-seq | MDA-MB-231 | 58111 | GSM1619769 | [61] |
| H3K9me3 | ChIP-seq | MDA-MB-468 | 87019 | GSM2258898 | [56] |
| H3K27me3 | ChIP-seq | HCC-1954 | 88398 | GSM2258832 | [56] |
| H3K27me3 | ChIP-seq | SKBR3 | 87171 | GSM2258796 | [56] |
| H3K27me3 | ChIP-seq | AU565 | 88150 | GSM2258814 | [56] |
| H3K27me3 | ChIP-seq | MDA-MB-361 | 81936 | GSM2258761 | [56] |
| H3K27me3 | ChIP-seq | MCF7 | 82436 | GSM2258724 | [56] |
| H3K27me3 | ChIP-seq | MDA-MB-231 | 88660 | GSM2258850 | [56] |
| H3K27me3 | ChIP-seq | MDA-MB-468 | 82827 | GSM2258887 | [56] |

### 4D genome data

IM-PET promoter-enhancer interaction data were obtained from 4Dgenome database [28] available at https://4dgenome.research.chop.edu.

### Statistical analysis and data visualization

GEO array expression data were analyzed by Affymetrix Transcriptome Analysis Console (TAC) 3.0 software (Affymetrix Inc., Santa Clara, CA, USA). Circus plots was created by Circa software respectively. Word could diagrams were generated using https://wordart.com web tool. All figure layouts were prepared using Adobe Photoshop CS6 (San Jose, CA, USA). Data were statistically analyzed by two-tailed student’s t-test and analysis of variance (ANOVA) using Prism v.6 software (GraphPad Software, La Jolla, CA, USA). Data were presented as mean and SD. *P* < 0.050 was considered as statistically significant.

## Acknowledgments

Our research was supported in part by a grant from Canadian Institutes of Health Research (CIHR) to Z.W. B.N. is supported by a Graduate Studentship from Women and Children’s Health Research Institute (WCHRI). We are thankful to CIHR and WCHRI for financial supports. No specific funding was received for this study.

## Author Contributions

All authors have read and approved the manuscript. B.N. participated in the design of the project, genomic data mining and analysis, statistical analysis, performance of all the experimental assays and manuscript writing. A.G. participated in genomic data mining and analysis. Z.W. participated in the design of the project, and in the data analysis and the writing of the manuscript.

## REFERENCES

1. Schechter, A.L.; Stern, D.F.; Vaidyanathan, L.; Decker, S.J.; Drebin, J.A.; Greene, M.I.; Weinberg, R.A. The neu oncogene: an erb-B-related gene encoding a 185,000-Mr tumour antigen. Nature 1984, 312, 513–516.

2. Slamon, D.J.; Godolphin, W.; Jones, L.A.; Holt, J.A.; Wong, S.G.; Keith, D.E.; Levin, W.J.; Stuart, S.G.; Udove, J.; Ullrich, A. Studies of the HER-2/neu proto-oncogene in human breast and ovarian cancer. Science 1989, 244, 707–712.

3. Carpenter, G. Receptors for epidermal growth factor and other polypeptide mitogens. Annu. Rev. Biochem. 1987, 56, 881–914.

4. Cho, H.-S.; Mason, K.; Ramyar, K.X.; Stanley, A.M.; Gabelli, S.B.; Denney, D.W.J.; Leahy, D.J. Structure of the extracellular region of HER2 alone and in complex with the Herceptin Fab. Nature 2003, 421, 756–760.

5. Aertgeerts, K.; Skene, R.; Yano, J.; Sang, B.-C.; Zou, H.; Snell, G.; Jennings, A.; Iwamoto, K.; Habuka, N.; Hirokawa, A.; et al. Structural analysis of the mechanism of inhibition and allosteric activation of the kinase domain of HER2 protein. J. Biol. Chem. 2011, 286, 18756–18765.

6. Garrett, T.P.J.; McKern, N.M.; Lou, M.; Elleman, T.C.; Adams, T.E.; Lovrecz, G.O.; Kofler, M.; Jorissen, R.N.; Nice, E.C.; Burgess, A.W.; et al. The crystal structure of a truncated ErbB2 ectodomain reveals an active conformation, poised to interact with other ErbB receptors. Mol. Cell 2003, 11, 495–505.

7. Pegram, M.D.; Konecny, G.; Slamon, D.J. The molecular and cellular biology of HER2/neu gene amplification/overexpression and the clinical development of herceptin (trastuzumab) therapy for breast cancer. Cancer Treat. Res. 2000, 103, 57–75.

8. Slamon, D.J.; Clark, G.M.; Wong, S.G.; Levin, W.J.; Ullrich, A.; McGuire, W.L. Human breast cancer: correlation of relapse and survival with amplification of the HER-2/neu oncogene. Science 1987, 235, 177–182.

9. Tandon, A.K.; Clark, G.M.; Chamness, G.C.; Ullrich, A.; McGuire, W.L. HER-2/neu oncogene protein and prognosis in breast cancer. J. Clin. Oncol. 1989, 7, 1120–1128.

10. Yarden, Y.; Sliwkowski, M.X. Untangling the ErbB signalling network. Nat. Rev. Mol. Cell Biol. 2001, 2, 127–137.

11. Nami, B.; Maadi, H.; Wang, Z. Mechanisms Underlying the Action and Synergism of Trastuzumab and Pertuzumab in Targeting HER2-Positive Breast Cancer. Cancers (Basel). 2018, 10.

12. Nami, B.; Wang, Z. HER2 in Breast Cancer Stemness: A Negative Feedback Loop towards Trastuzumab Resistance. Cancers (Basel). 2017, 9.

13. Nami, B.; Maadi, H.; Wang, Z. The Effects of Pertuzumab and Its Combination with Trastuzumab on HER2 Homodimerization and Phosphorylation. Cancers (Basel). 2019, 11.

14. Maadi, H.; Nami, B.; Tong, J.; Li, G.; Wang, Z. The effects of trastuzumab on HER2-mediated cell signaling in CHO cells expressing human HER2. BMC Cancer 2018, 18, 238.

15. Ahmed, S.; Sami, A.; Xiang, J. HER2-directed therapy: current treatment options for HER2-positive breast cancer. Breast Cancer 2015, 22, 101–116.

16. Tural, D.; Akar, E.; Mutlu, H.; Kilickap, S. P95 HER2 fragments and breast cancer outcome. Expert Rev. Anticancer Ther. 2014, 14, 1089–1096.

17. Arribas, J.; Baselga, J.; Pedersen, K.; Parra-Palau, J.L. p95HER2 and breast cancer. Cancer Res. 2011, 71, 1515–1519.

18. Jeon, M.; Lee, J.; Nam, S.J.; Shin, I.; Lee, J.E.; Kim, S. Induction of fibronectin by HER2 overexpression triggers adhesion and invasion of breast cancer cells. Exp. Cell Res. 2015, 333, 116–126.

19. Oliveras-Ferraros, C.; Vazquez-Martin, A.; Martin-Castillo, B.; Cufi, S.; Del Barco, S.; Lopez-Bonet, E.; Brunet, J.; Menendez, J.A. Dynamic emergence of the mesenchymal CD44(pos)CD24(neg/low) phenotype in HER2-gene amplified breast cancer cells with de novo resistance to trastuzumab (Herceptin). Biochem. Biophys. Res. Commun. 2010, 397, 27–33.

20. Reim, F.; Dombrowski, Y.; Ritter, C.; Buttmann, M.; Hausler, S.; Ossadnik, M.; Krockenberger, M.; Beier, D.; Beier, C.P.; Dietl, J.; et al. Immunoselection of breast and ovarian cancer cells with trastuzumab and natural killer cells: selective escape of CD44high/CD24low/HER2low breast cancer stem cells. Cancer Res. 2009, 69, 8058–8066.

21. Pereira, B.; Chin, S.-F.; Rueda, O.M.; Vollan, H.-K.M.; Provenzano, E.; Bardwell, H.A.; Pugh, M.; Jones, L.; Russell, R.; Sammut, S.-J.; et al. The somatic mutation profiles of 2,433 breast cancers refines their genomic and transcriptomic landscapes. Nat. Commun. 2016, 7, 11479.

22. Cerami, E.; Gao, J.; Dogrusoz, U.; Gross, B.E.; Sumer, S.O.; Aksoy, B.A.; Jacobsen, A.; Byrne, C.J.; Heuer, M.L.; Larsson, E.; et al. The cBio cancer genomics portal: an open platform for exploring multidimensional cancer genomics data. Cancer Discov. 2012, 2, 401–404.

23. Dezso, Z.; Oestreicher, J.; Weaver, A.; Santiago, S.; Agoulnik, S.; Chow, J.; Oda, Y.; Funahashi, Y. Gene expression profiling reveals epithelial mesenchymal transition (EMT) genes can selectively differentiate eribulin sensitive breast cancer cells. PLoS One 2014, 9, e106131.

24. Xiao, Y.; Nimmer, P.; Sheppard, G.S.; Bruncko, M.; Hessler, P.; Lu, X.; Roberts-Rapp, L.; Pappano, W.N.; Elmore, S.W.; Souers, A.J.; et al. MCL-1 Is a Key Determinant of Breast Cancer Cell Survival: Validation of MCL-1 Dependency Utilizing a Highly Selective Small Molecule Inhibitor. Mol. Cancer Ther. 2015, 14, 1837–1847.

25. Di Cello, F.; Cope, L.; Li, H.; Jeschke, J.; Wang, W.; Baylin, S.B.; Zahnow, C.A. Methylation of the claudin 1 promoter is associated with loss of expression in estrogen receptor positive breast cancer. PLoS One 2013, 8, e68630.

26. Mei, S.; Qin, Q.; Wu, Q.; Sun, H.; Zheng, R.; Zang, C.; Zhu, M.; Wu, J.; Shi, X.; Taing, L.; et al. Cistrome Data Browser: a data portal for ChIP-Seq and chromatin accessibility data in human and mouse. Nucleic Acids Res. 2017, 45, D658–62.

27. Zhou, X.; Lowdon, R.F.; Li, D.; Lawson, H.A.; Madden, P.A.F.; Costello, J.F.; Wang, T. Exploring long-range genome interactions using the WashU Epigenome Browser. Nat. Methods 2013, 10, 375–376.

28. Teng, L.; He, B.; Wang, J.; Tan, K. 4DGenome: a comprehensive database of chromatin interactions. Bioinformatics 2015, 31, 2560–2564.

29. Liu, L.; Greger, J.; Shi, H.; Liu, Y.; Greshock, J.; Annan, R.; Halsey, W.; Sathe, G.M.; Martin, A.-M.; Gilmer, T.M. Novel mechanism of lapatinib resistance in HER2-positive breast tumor cells: activation of AXL. Cancer Res. 2009, 69, 6871–6878.

30. Sartor, M.A.; Mahavisno, V.; Keshamouni, V.G.; Cavalcoli, J.; Wright, Z.; Karnovsky, A.; Kuick, R.; Jagadish, H. V; Mirel, B.; Weymouth, T.; et al. ConceptGen: a gene set enrichment and gene set relation mapping tool. Bioinformatics 2010, 26, 456–463.

31. Liu, H.; Patel, M.R.; Prescher, J.A.; Patsialou, A.; Qian, D.; Lin, J.; Wen, S.; Chang, Y.-F.; Bachmann, M.H.; Shimono, Y.; et al. Cancer stem cells from human breast tumors are involved in spontaneous metastases in orthotopic mouse models. Proc. Natl. Acad. Sci. U. S. A. 2010, 107, 18115–18120.

32. Mani, S.A.; Guo, W.; Liao, M.-J.; Eaton, E.N.; Ayyanan, A.; Zhou, A.Y.; Brooks, M.; Reinhard, F.; Zhang, C.C.; Shipitsin, M.; et al. The epithelial-mesenchymal transition generates cells with properties of stem cells. Cell 2008, 133, 704–715.

33. Giordano, A.; Mego, M.; Lee, B.; Anfossi, S.; Parker, C.A.; Alvarez, R.H.; Ueno, N.T.; Valero, V.; Cristofanilli, M.; Reuben, J.M. Epithelial-mesenchymal transition in patients with HER2+ metastatic breast cancer. J. Clin. Oncol. 2011, 29, 623.

34. Giordano, A.; Gao, H.; Anfossi, S.; Cohen, E.; Mego, M.; Lee, B.-N.; Tin, S.; De Laurentiis, M.; Parker, C.A.; Alvarez, R.H.; et al. Epithelial-mesenchymal transition and stem cell markers in patients with HER2-positive metastatic breast cancer. Mol. Cancer Ther. 2012, 11, 2526–2534.

35. Jenndahl, L.E.; Isakson, P.; Baeckstrom, D. c-erbB2-induced epithelial-mesenchymal transition in mammary epithelial cells is suppressed by cell-cell contact and initiated prior to E-cadherin downregulation. Int. J. Oncol. 2005, 27, 439–448.

36. Nilsson, G.M.A.; Akhtar, N.; Kannius-Janson, M.; Baeckstrom, D. Loss of E-cadherin expression is not a prerequisite for c-erbB2-induced epithelial-mesenchymal transition. Int. J. Oncol. 2014, 45, 82–94.

37. Ingthorsson, S.; Andersen, K.; Hilmarsdottir, B.; Maelandsmo, G.M.; Magnusson, M.K.; Gudjonsson, T. HER2 induced EMT and tumorigenicity in breast epithelial progenitor cells is inhibited by coexpression of EGFR. Oncogene 2016, 35, 4244–4255.

38. Rennstam, K.; Jonsson, G.; Tanner, M.; Bendahl, P.-O.; Staaf, J.; Kapanen, A.I.; Karhu, R.; Baldetorp, B.; Borg, A.; Isola, J. Cytogenetic characterization and gene expression profiling of the trastuzumab-resistant breast cancer cell line JIMT-1. Cancer Genet. Cytogenet. 2007, 172, 95–106.

39. Barok, M.; Isola, J.; Palyi-Krekk, Z.; Nagy, P.; Juhasz, I.; Vereb, G.; Kauraniemi, P.; Kapanen, A.; Tanner, M.; Vereb, G.; et al. Trastuzumab causes antibody-dependent cellular cytotoxicity-mediated growth inhibition of submacroscopic JIMT-1 breast cancer xenografts despite intrinsic drug resistance. Mol. Cancer Ther. 2007, 6, 2065–2072.

40. Koninki, K.; Barok, M.; Tanner, M.; Staff, S.; Pitkanen, J.; Hemmila, P.; Ilvesaro, J.; Isola, J. Multiple molecular mechanisms underlying trastuzumab and lapatinib resistance in JIMT-1 breast cancer cells. Cancer Lett. 2010, 294, 211–219.

41. Cavaliere, C.; Corvigno, S.; Galgani, M.; Limite, G.; Nardone, A.; Veneziani, B.M. Combined inhibitory effect of formestane and herceptin on a subpopulation of CD44+/CD24low breast cancer cells. Cancer Sci. 2010, 101, 1661–1669.

42. Oliveras-Ferraros, C.; Corominas-Faja, B.; Cufi, S.; Vazquez-Martin, A.; Martin-Castillo, B.; Iglesias, J.M.; Lopez-Bonet, E.; Martin, A.G.; Menendez, J.A. Epithelial-to-mesenchymal transition (EMT) confers primary resistance to trastuzumab (Herceptin). Cell Cycle 2012, 11, 4020–4032.

43. Cufí, S.; Corominas-Faja, B.; Vazquez-Martin, A.; Oliveras-Ferraros, C.; Dorca, J.; Bosch-Barrera, J.; Martin-Castillo, B.; Menendez, J.A. Metformin-induced preferential killing of breast cancer initiating CD44(+)CD24(-/low) cells is sufficient to overcome primary resistance to trastuzumab in HER2+ human breast cancer xenografts. Oncotarget 2012, 3, 395–398.

44. Bauerschmitz, G.J.; Ranki, T.; Kangasniemi, L.; Ribacka, C.; Eriksson, M.; Porten, M.; Herrmann, I.; Ristimaki, A.; Virkkunen, P.; Tarkkanen, M.; et al. Tissue-specific promoters active in CD44+CD24-/low breast cancer cells. Cancer Res. 2008, 68, 5533–5539.

45. Rennstam, K.; McMichael, N.; Berglund, P.; Honeth, G.; Hegardt, C.; Ryden, L.; Luts, L.; Bendahl, P.-O.; Hedenfalk, I. Numb protein expression correlates with a basal-like phenotype and cancer stem cell markers in primary breast cancer. Breast Cancer Res. Treat. 2010, 122, 315–324.

46. Lesniak, D.; Sabri, S.; Xu, Y.; Graham, K.; Bhatnagar, P.; Suresh, M.; Abdulkarim, B. Spontaneous Epithelial-Mesenchymal Transition and Resistance to HER-2-Targeted Therapies in HER-2-Positive Luminal Breast Cancer. PLoS One 2013, 8.

47. Nami, B.; Wang, Z. Application of Immunofluorescence Staining to Study ErbB Family of Receptor Tyrosine Kinases. Methods Mol. Biol. 2017, 1652, 109–116.

48. Gao, J.; Aksoy, B.A.; Dogrusoz, U.; Dresdner, G.; Gross, B.; Sumer, S.O.; Sun, Y.; Jacobsen, A.; Sinha, R.; Larsson, E.; et al. Integrative analysis of complex cancer genomics and clinical profiles using the cBioPortal. Sci. Signal. 2013, 6, pl1.

49. Porter, J.R.; Fisher, B.E.; Baranello, L.; Liu, J.C.; Kambach, D.M.; Nie, Z.; Koh, W.S.; Luo, J.; Stommel, J.M.; Levens, D.; et al. Global Inhibition with Specific Activation: How p53 and MYC Redistribute the Transcriptome in the DNA Double-Strand Break Response. Mol. Cell 2017, 67, 1013–1025.e9.

50. Bustos, M.A.; Salomon, M.P.; Nelson, N.; Hsu, S.C.; DiNome, M.L.; Hoon, D.S.B.; Marzese, D.M. Genome-wide chromatin accessibility, DNA methylation and gene expression analysis of histone deacetylase inhibition in triple-negative breast cancer. Genomics data 2017, 12, 14–16.

51. Thurman, R.E.; Rynes, E.; Humbert, R.; Vierstra, J.; Maurano, M.T.; Haugen, E.; Sheffield, N.C.; Stergachis, A.B.; Wang, H.; Vernot, B.; et al. The accessible chromatin landscape of the human genome. Nature 2012, 489, 75–82.

52. Gallenne, T.; Ross, K.N.; Visser, N.L.; Salony; Desmet, C.J.; Wittner, B.S.; Wessels, L.F.A.; Ramaswamy, S.; Peeper, D.S. Systematic functional perturbations uncover a prognostic genetic network driving human breast cancer. Oncotarget 2017, 8, 20572–20587.

53. Ni, M.; Chen, Y.; Fei, T.; Li, D.; Lim, E.; Liu, X.S.; Brown, M. Amplitude modulation of androgen signaling by c-MYC. Genes Dev. 2013, 27, 734–748.

54. Joseph, R.; Orlov, Y.L.; Huss, M.; Sun, W.; Kong, S.L.; Ukil, L.; Pan, Y.F.; Li, G.; Lim, M.; Thomsen, J.S.; et al. Integrative model of genomic factors for determining binding site selection by estrogen receptor-alpha. Mol. Syst. Biol. 2010, 6, 456.

55. Cao, A.R.; Rabinovich, R.; Xu, M.; Xu, X.; Jin, V.X.; Farnham, P.J. Genome-wide analysis of transcription factor E2F1 mutant proteins reveals that N- and C-terminal protein interaction domains do not participate in targeting E2F1 to the human genome. J. Biol. Chem. 2011, 286, 11985–11996.

56. Franco, H.L.; Nagari, A.; Malladi, V.S.; Li, W.; Xi, Y.; Richardson, D.; Allton, K.L.; Tanaka, K.; Li, J.; Murakami, S.; et al. Enhancer transcription reveals subtype-specific gene expression programs controlling breast cancer pathogenesis. Genome Res. 2018, 28, 159–170.

57. Hodges, E.; Smith, A.D.; Kendall, J.; Xuan, Z.; Ravi, K.; Rooks, M.; Zhang, M.Q.; Ye, K.; Bhattacharjee, A.; Brizuela, L.; et al. High definition profiling of mammalian DNA methylation by array capture and single molecule bisulfite sequencing. Genome Res. 2009, 19, 1593–1605.

58. Hon, G.C.; Hawkins, R.D.; Caballero, O.L.; Lo, C.; Lister, R.; Pelizzola, M.; Valsesia, A.; Ye, Z.; Kuan, S.; Edsall, L.E.; et al. Global DNA hypomethylation coupled to repressive chromatin domain formation and gene silencing in breast cancer. Genome Res. 2012, 22, 246–258.

59. Messier, T.L.; Gordon, J.A.R.; Boyd, J.R.; Tye, C.E.; Browne, G.; Stein, J.L.; Lian, J.B.; Stein, G.S. Histone H3 lysine 4 acetylation and methylation dynamics define breast cancer subtypes. Oncotarget 2016, 7, 5094–5109.

60. Zhu, J.; Sammons, M.A.; Donahue, G.; Dou, Z.; Vedadi, M.; Getlik, M.; Barsyte-Lovejoy, D.; Al-awar, R.; Katona, B.W.; Shilatifard, A.; et al. Gain-of-function p53 mutants co-opt chromatin pathways to drive cancer growth. Nature 2015, 525, 206–211.

61. Vasudevan, D.; Hickok, J.R.; Bovee, R.C.; Pham, V.; Mantell, L.L.; Bahroos, N.; Kanabar, P.; Cao, X.-J.; Maienschein-Cline, M.; Garcia, B.A.; et al. Nitric Oxide Regulates Gene Expression in Cancers by Controlling Histone Posttranslational Modifications. Cancer Res. 2015, 75, 5299–5308.

62. Takaku, M.; Grimm, S.A.; Shimbo, T.; Perera, L.; Menafra, R.; Stunnenberg, H.G.; Archer, T.K.; Machida, S.; Kurumizaka, H.; Wade, P.A. GATA3-dependent cellular reprogramming requires activation-domain dependent recruitment of a chromatin remodeler. Genome Biol. 2016, 17, 36.

